# ZnF-UBP domains regulate deubiquitinase activity by relieving ubiquitin product inhibition

**DOI:** 10.1101/2025.09.28.679104

**Authors:** Jack A. Alexandrovics, Rashmi Agrata, Philipp Schenk, Anthony Cerra, Ueli Nachbur, Jeffrey J. Babon, David Komander

## Abstract

Most ubiquitin specific protease (USP) deubiquitinases (DUBs) combine non-selective catalytic domains with one or multiple ‘exo’-domains that contribute substrate specificity and localisation, but are generally poorly characterised. Zinc-Finger UBP (ZnF-UBP) domains exist in 12 USP DUBs, yet their function is unclear.

We here comprehensively analyse human ZnF-UBP domains, and reveal that 8 of 14 bind ubiquitin (Ub), via an unattached Ub C-terminal GlyGly motif. We focus on USP16, a nucleosome DUB with activity for Ub and Ub-like modifiers, and show that whilte its ZnF-UBP domain can bind substrates, it is also a crucial contributor to enzyme kinetics. Slow Ub release from the catalytic domain after cleavage causes product inhibition, which is overcome in *cis* by ZnF-UBP-mediated product release. Interestingly, supplying a high affinity product-capturing ZnF-UBP domain in *trans*, activates USP16 and other USP enzymes. Our data shows the importance of product inhibition as a regulatory mechanism in DUBs, and exemplifies the unappreciated role of exo-domains in regulating DUB function beyond substrate binding.

## Introduction

Protein modifications by the small protein ubiquitin (Ub) comprise a complex Ub code^1^ that controls cellular processes prolifically and in unrivalled breadth. Ubiquitination typically targets Lys residues in protein substrates, which is facilitated by a sophisticated enzymatic assembly cascade^1^. The Ub code is read by Ub binding domains (UBDs)^2^ found across many proteins, which recognise accessible surfaces on Ub or on specific Ub chain topologies^3^. Deubiquitinases (DUBs) are enzymes responsible for removing Ub modifications. Approximately 100 human DUBs across seven families read, reverse or fine-tune Ub signalling, with fascinating mechanistic variety^4,5^. While some DUB families regulate specific signals, the 56 members of the ubiquitin specific protease (USP) family feature catalytic domains that, typically, remove Ub modifications without specificity^4^. Some USPs attain functional specificity through incorporation into complexes^6^, yet the majority achieve it through ‘exo’ domains that extend beyond the ∼350 amino acid (aa) catalytic domain, creating some of the largest enzymes in the human genome^5,7,8^.

UBDs are highly represented as exo-domains in DUBs and enable the enzyme to select Ub signals on substrates^9,10^. USP5 is a highly abundant DUB with the important role to cleave unattached Ub chains, generated from polyUb genes or by endo-DUB activity, into ‘free’ monoUb units^11^. A UBD in USP5, the ZnF-UBP domain, specifically recognises Ub by its unconjugated C-terminal GlyGly motif^12^. The ZnF-UBP domain hence binds the proximal Ub in free polyUb, anchoring the chain and positioning the adjacent, more distal Ub molecules, for cleavage by the catalytic domain^10,13^. Strikingly, in the human genome there are 12 USPs as well as two non-DUBs (the histone deacetylase HDAC6 and the E3 ligase BRAP2) that contain ZnF-UBP domains (**Fig. 1a**)^14,15^. Structures for USP5^12^ and HDAC6 ZnF-UBPs^16^ in complex with Ub revealed a shared recognition mode for the Ub C-terminal tail and the GlyGly motif, yet distinct binding orientations for the Ub fold (**Fig. 1b**). Conversely, the ZnF-UBP domains of USP22^17,18^ and USP39^19^ act as structural components within the SAGA (Spt-Ada-Gcn5 acetyltransferase) complex and the spliceosome, respectively. While several other ZnF-UBP domains have been biochemically characterised^16,20–24^, a clear picture for shared functionality or role has not emerged. Indeed, the binding specificity of ZnF-UBP domains to Ub, their dependence on free Ub, and the underlying reasons for this unexpected interaction remain unclear.

**Figure 1.**
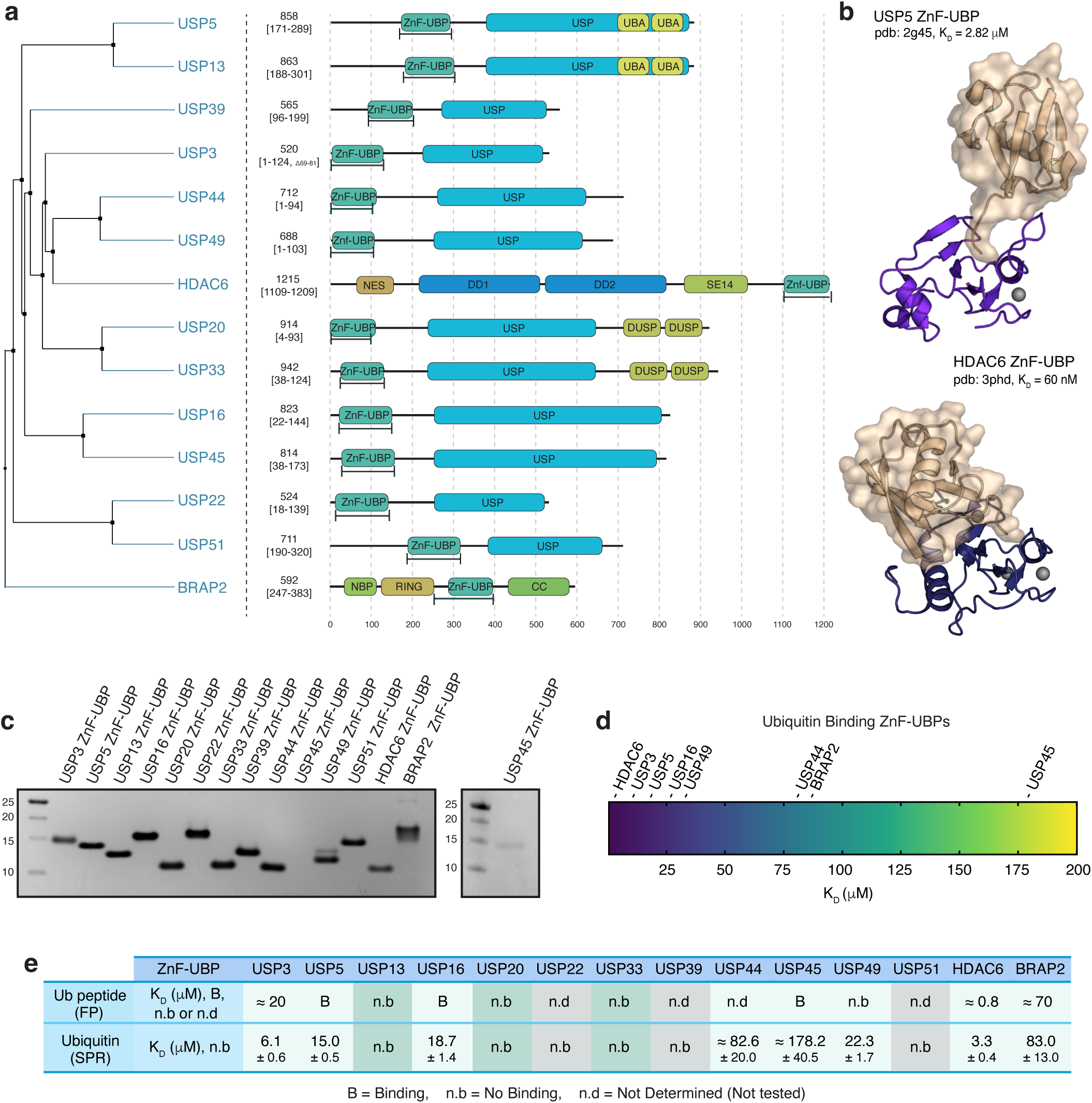
Characterising Ub binding in ZnF-UBP domains. **a,** Phylogenetic tree of human ZnF-UBP domains and domain organization of the full-length ZnF-UBP-containing enzymes. Expressed ZnF-UBP construct boundaries are shown in square brackets. **b,** Available ZnF-UBP:Ub complex structures for USP5 (2g45) and HDAC6 (3phd) and their reported binding affinities^12,70^. 1ubq was used to complete missing density in 3phd. **c,** Coomassie stained SDS-PAGE gels of purified ZnF-UBP domains corresponding to the construct boundaries in **a**. USP45 ZnF-UBP expressed at low yields and is visualised on a separate gel. **d,** Heatmap of relative binding affinities for the Ub-binding ZnF-UBP domains as determined by SPR. Also see **Extended Data Fig. 1**. **e,** Comprehensive summary of ZnF-UBP Ub-binding data with affinity determined by FP and SPR. When full binding curves could not be achieved in FP due to concentration limitations, results are reported as Binding (B) or non-binding (n.b). For raw data see **Extended Data Fig. 1**. Errors correspond to standard deviation (s.d.) from the mean.

Seven ZnF-UBP containing USPs (USP3, USP16, USP22, USP27X, USP44, USP49, USP51) reportedly deubiquitinate histones^20,25–34^, which constitute the most abundant source of monoubiquitination in cells. Nucleosome ubiquitination is an epigenetic mark that regulates gene expression^33^. The listed DUBs contain a ZnF-UBP domain as their sole exo-domain (**Fig. 1a**) yet the role of the ZnF-UBP domain has been minimally explored, with exception of USP22, in which case the ZnF-UBP domain is an integral scaffolding part of the SAGA complex^17,18^. A recent structure resolved USP16 in complex with nucleosome H2AK119Ub^35^, however whilst present in the studied protein, the ZnF-UBP domain was not resolved in the electron density, and was not discussed further. The cellular functions of USP16 (previously UBP-M^36^) appear pleiotropic, acting on a variety of substrates, chain types, and even Ub-like proteins such as Fubi^37–40^. USP16’s nucleosome deubiquitination has been implicated in cell cycle regulation during development and has been associated with developmental disorders^41–45^, with cell biology studies interrogating its pathways and process of nuclear localization^46–49^. Notably, the ZnF-UBP domain of USP16 both binds Ub and appears to havea role in nucleosome interactions, despite its absence in the nucleosome structure^22,50–52^ yet an overall mechanism for this DUB remains elusive. Beyond USP16, this raises intriguing questions about the purpose of Ub binding ZnF-UBP domains existing on histone DUBs, given the known yet unclear links between Ub pool dynamics, proteostasis and gene regulation^53–55^.

Indeed, ZnF-UBP domains acting as ‘sensors’ of free Ub has been suggested^15,22,56^, and some DUBs are already known to respond to Ub pool dynamics^13,57–60^. Any effects from the Ub pool are intrinsically linked to Ub concentration^61–63^ which in turn affects affinity and cleavage kinetics, parameters often governed by UBDs in DUBs^4,38^. Ablation of ZnF-UBP Ub binding in USP5 affected kinetic parameters and Ub interactions, and similar effects have been seen for UBDs across DUBs, including USP16^38^. This highlights the need to understand the broader and complex role of Ub interactions and kinetics in shaping DUB activity^64,65^, within the highly dynamic and Ub-rich environment of the cell.

We here provide a comprehensive characterisation of all human ZnF-UBP domains, cross-comparing Ub binding capabilities, and structurally explaining our findings. We then focus on USP16, and show that it binds the histone H4 tail with low affinity, but is also an integral part of enzyme mechanism, and dislodges cleaved Ub from the catalytic domain.

## Results

### ZnF UBP domains show a varied profile of ubiquitin interactions

Current work on ZnF-UBP domains is fragmented, but data suggests not all bind Ub, and that Ub interactions are facilitated with varied mechanisms and affinities. We hence annotated (**Fig. 1a**) and expressed the ZnF-UBP domains of 14 human enzymes in *E. coli* (**Fig. 1c**). Most domains expressed well but yields for USP44, USP51 and especially USP45 ZnF-UBP were low. Not annotated genetically, USP27X appears to express with a preceding ZnF-UBP domain from an unconventional start codon and acts akin USP22 to provide scaffolding within the SAGA complex^66^; as this ZnF-UBP originated from alternative splicing we did not include it in this study. We assessed Ub binding capabilities with two independent assays. We used fluorescence polarisation (FP) to study the ZnF-UBP interaction with a fluorescently labelled Ub tail peptide (FITC-Ahx-RLRGG, see **Methods**, hereafter UbCt peptide) (**Extended Data Fig. 1a**, **Fig. 1e**). Similar assays have been performed previously, and we largely replicated published results for USP16 and HDAC6^51,67^, which were in a similar range of affinities compared to other UbCt-peptide based studies^68,69^. However, in FP studies only HDAC6 interacted with the UbCt-peptide with high enough affinity to fit a binding curve; USP3, USP5, USP16, USP45 and BRAP2 showed clear but low affinity (**Extended Data Fig. 1a**, **Fig. 1e**).

The ZnF-UBP domain of USP5 binds not only the Ub C-terminus but also makes secondary contacts (**Fig. 1b**). We hence performed Surface Plasmon Resonance (SPR) experiments with immobilised Ub. Similarly to the FP assay, no binding was detected for USP13, USP20, USP22, USP33 and USP39, while affinities between 6 and ∼175 µM were measured for the remaining ZnF-UBP domains (**Fig. 1d, e, Extended Data Fig. 1b**). USP49 was interesting since it interacted strongly with Ub in SPR, but did not interact with the UbCt peptide, suggesting additional binding modes (**Extended Data Fig. 1**, **Fig. 1e**). Our data largely aligns with published reports^12,22,70^, revealing that 8 of 14 human ZnF-UBP domains bind Ub, and 7 bind via the unattached UbCt peptide. Five Ub affinities are sub-25 µM, which is considered high affinity for Ub interactions^4^. The 6 ZnF-UBP domains that do not bind Ub are all in USP enzymes and may have structural or substrate binding roles.

### Structural basis for ZnF-UBP ubiquitin interactions

To understand the molecular basis for Ub interactions in DUB ZnF-UBPs, we considered sequence requirements (**Extended Data Fig. 2a**) and structural features (**Fig. 2a**). Several structures are available, including Ub complex structures (**Fig. 1b**), isolated ZnF-UBP NMR structures^22–24^, and as part of large macromolecular assemblies^18,19^ (**Fig. 2a**). We here report crystal structures of Ub-binding ZnF-UBPs USP16 at 1.8 Å, and of USP49 at 1.41 Å resolution (**Fig. 2a-c, Extended Data Fig. 2b, Extended Data Table 1**). The USP16 ZnF-UBP asymmetric unit contains three ZnF-UBP molecules which overlap with an RMSD of <0.25 Å, whereas the USP49 ZnF-UBP asymmetric unit contains a single molecule (**Extended Data Fig. 2b**). Each ZnF-UBP domain coordinates three Zn-ions, and both domains closely overlap with AlphaFold3 predictions. AlphaFold3 was used to model the remaining domains with high confidence (**Fig. 2a**).

**Figure 2.**
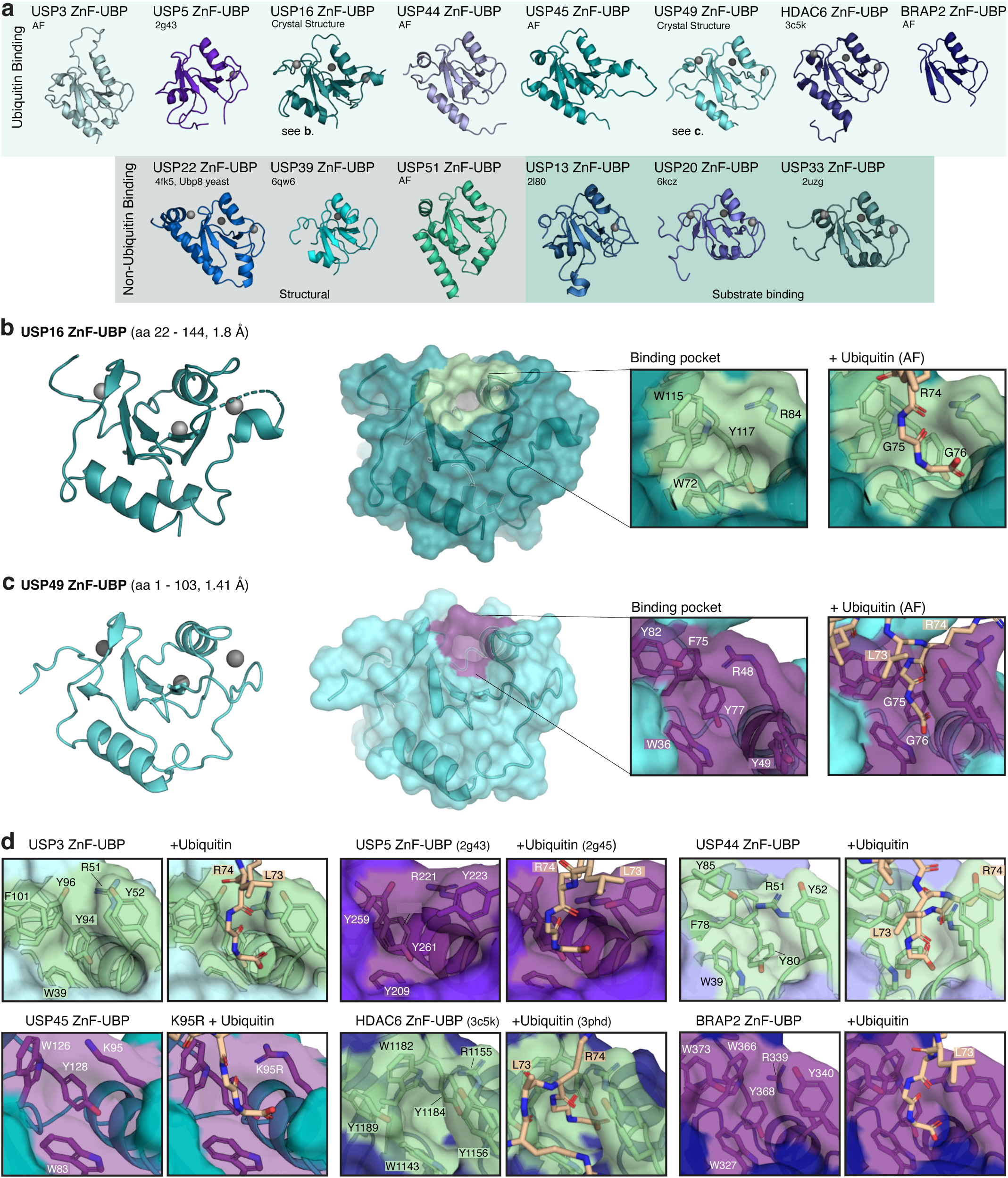
Structural basis for Ub interactions. **a,** Experimental and AlphaFold3-predicted structures of ZnF-UBP domains grouped according to their Ub-binding capacity. Non-Ub binding domains are grouped based on known structural roles or proposed functions in substrate interaction^83,84^. Shading correlates to Fig. 1e. **b,** Crystal structure of the USP16 ZnF-UBP domain shown in cartoon (*left*) and surface (*centre*) representations. The Ub binding pocket is coloured pale green, with close-up views highlighting binding residues in apo form (*left*) and with Ub tail residues from an Ub-bound AlphaFold3-modelled prediction (*right*). See **Extended Data Fig. 2b** and **Extended Data Table 1**. **d,** Crystal structure of USP49 ZnF-UBP domain with the binding pocket coloured purple, and visualised as in **c**. **d,** Close-up views of ZnF-UBP binding pockets, highlighting the ZnF-UBP binding residues (*left*) and the Ub residues (*right*; G75 and G76 labels omitted for clarity). The K95R mutation is labelled for USP45, and was required for AlphaFold3 to predict a Ub-bound model. All models were generated through AlphaFold3-predictions with or without Ub, except for USP5 (2g45) and HDAC6 (3phd). Leu73 was modelled as an Ala in 3phd.

All Ub-binding ZnF-UBPs show prominent pockets poised to interact with the Ub C-terminal tail, and AlphaFold3 predicted these binding interfaces confidently (**Fig. 2d**). In contrast, non-Ub-binding ZnF-UBP domains lack the UbCt peptide pocket, and Ub complex modelling does not place Ub in or near this surface (not shown). At the sequence level, the structures explain the role of a key Arg residue and a suite of up to 5 aromatic residues, in Ub-interactions (**Extended Data Fig. 2a**). As in USP5^12^, the key Arg guides Ub backbone carbonyls of Arg74 and Gly75 into the UbCt pocket, while the terminal Gly interacts with several hydrophobic pocket residues through the α carbon and carboxyl group. Modelling Ub binding for low affinity (**Fig. 1d**) USP45 placed Ub at a different surface, without involving the otherwise existing UbCt binding pocket, similar to modelling performed for other ZnF-UBPs^23^. Interestingly, USP45 contains a Lys instead of canonical Arg residue and when we modelled with USP45 K95R mutation, canonical Ub binding was predicted by AlphaFold3 (**Fig. 2d, Extended Data Fig. 2c**). Further insights from structural modelling are discussed in the **Supplementary Text**. Together, experimental structures and AlphaFold3 models enable a comprehensive molecular understanding of ZnF-UBP domain Ub binding mechanisms, which were exploited for functional studies below.

### Understanding the contributions of the ZnF-UBP domain in USP16

We next investigated whether and how the USP16 ZnF-UBP domain contributed its reported functions as a nucleosome DUB and as a deFubiylase^39^, and tested whether the canonical UbCt pocket in the USP16 ZnF-UBP domain could contribute to substrate interactions. Peptide binding to this domain has been comprehensively studied^51^, and both the histone H4 tail and Fubi had suitable C-terminal sequences (**Fig. 3a, Extended Data Fig. 3a**). We performed SPR and FP assays with Fubi and histone H4 peptides which showed clear binding of both to the ZnF-UBP domain, although with 3-fold (Fubi) and >10-fold (histone H4) weaker affinity as compared to the UbCt peptide (**Fig. 3a, Extended Data Fig. 3a**). AlphaFold3 modelling suggested that both peptides could interact with the UbCt, and residue differences showed where steric interactions might affect peptide entry (**Fig. 3a**). We validated the interaction with the histone H4 peptide using NMR (**Fig. 3b, Extended Data Fig. 3b, c**). For this, we generated ^15^N-labelled USP16 ZnF-UBP, and spectra were annotated based on the published assignments (BMRB_7298)^22^. ^15^N-labelled ZnF-UBP was titrated with unlabelled histone H4 peptide, which yielded chemical shift perturbations (CSPs) in a subset of resonances. The CSPs are indicative of interactions, and shared key similarities with perturbations observed with Ub and a UbCt peptide in a previous study^22^. The key Ub-binding residue Arg84 (**Fig. 3a**) was not unanimously assigned, but resonance of the adjacent Asn85 shifted markedly, with this trend extending nearby to Lys93 (**Fig. 3b**), consistent with involvement in peptide interaction^22^. The Ser86 peak disappeared upon binding suggesting strong interaction. Of the key Ub-binding pocket residues, only Tyr117 showed pronounced shifts, yet notably, residues lining the pocket, Asp120 to Glu122, were also perturbed. Asp264 in USP5, equivalent to Asp120 in USP16, has previously been shown to coordinate Ub tail interactions^12^, and the significant shifts in this region indicate a widening of the pocket to accommodate the more-bulky histone H4 peptide, corroborated by compression of Cys119 and Val123 (**Extended Data Fig. 3c**). Taken together, these data show that histone H4 binds weaker than Ub to USP16 ZnF-UBP domain, likely due to the bulky Phe101, located upstream of the histone H4 GlyGly terminus, preventing the peptide from penetrating deeply (**Fig. 3a, Extended Data Fig. 3c**).

**Figure 3.**
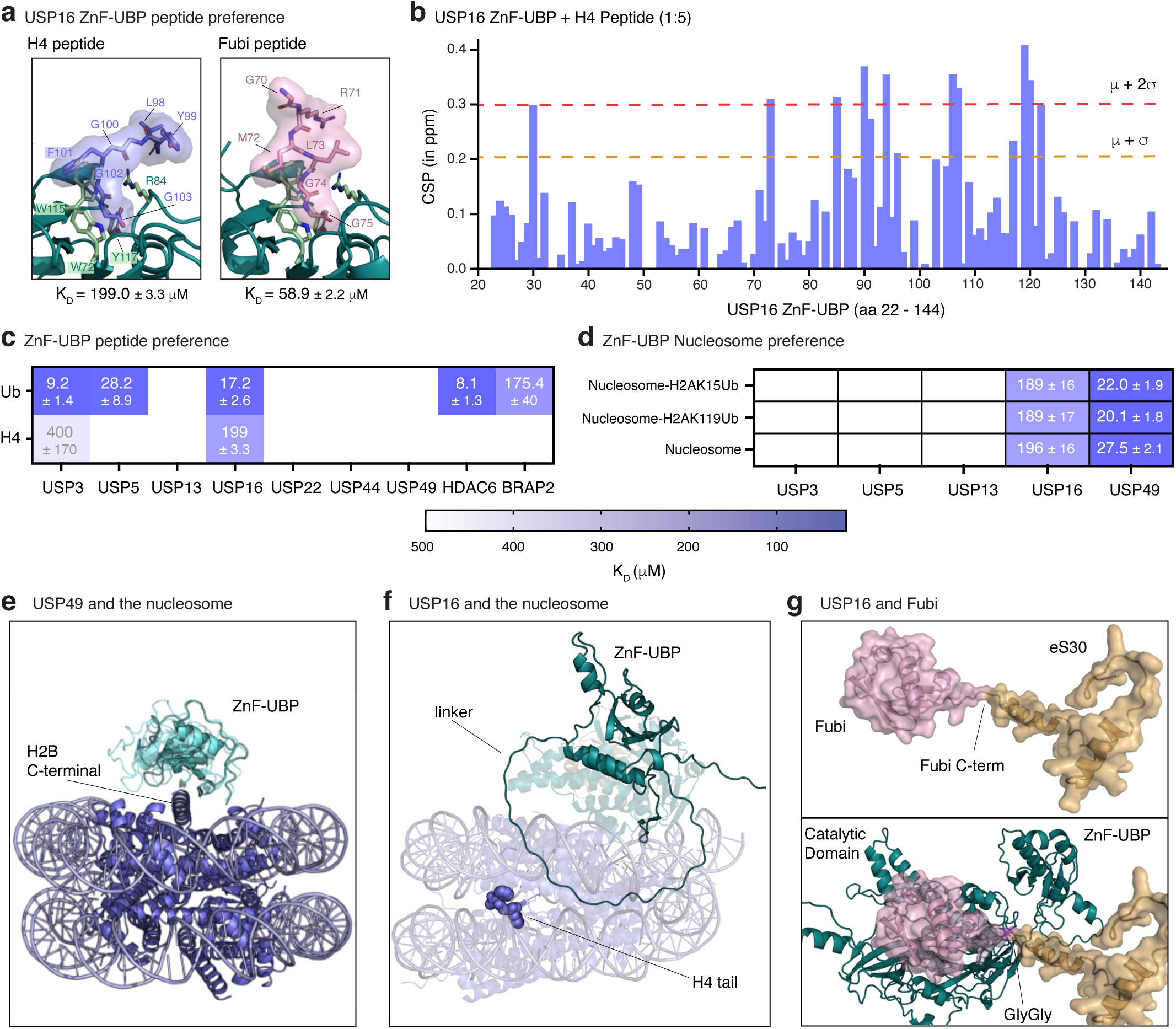
ZnF-UBP domains may regulate substrate interactions. **a,** AlphaFold3 prediction for USP16 ZnF-UBP with histone H4 tail peptide, and Fubi tail peptide. K_D_ values were derived from SPR data in **Extended Data Fig. 3a**. Errors correspond to s.d. from the mean. **b,** The CSPs of ¹⁵N-labelled USP16 ZnF-UBP upon H4 peptide binding (1:5 ratio) plotted against its residues. The red and the orange dotted lines correspond to mean+s.d. and mean+2*s.d., respectively **c,** Comparison of Ub and histone H4 tail peptide binding affinities in µM, to select ZnF-UBP domains, derived from SPR data in **Extended Data Fig. 3d**. Errors correspond to s.d. from the mean. **d,** Heatmap of binding affinities in µM, derived from SPR in **Extended Data Fig. 3e-f**, for indicated ZnF-UBP domains against indicated nucleosome variants. Errors correspond to s.d. from the mean. **e,** AlphaFold3-predicted model of USP49 ZnF-UBP binding to the H2B C-terminus in an unubiquitinated nucleosome. **f,** AlphaFold3-predicted model of full-length USP16 aligned to USP16:nucleosomeH2AK119Ub structure (pdb-id 8wg5). The histone H4 tail is shown as blue spheres, but is occluded between histones and DNA. **g,** AlphaFold3-model prediction of Fubi-eS30 apo (*top*) and bound to full-length USP16 (*bottom*).

We next tested whether other ZnF-UBP domains can bind the histone H4 tail. Interestingly, histone tail interaction appears to be restricted to USP16 and at even weaker affinity of 400 µM, to USP3 in SPR experiments (**Fig. 3c, Extended Data Fig. 3d**). Because peptide assays miss chromatin-dependent contacts, we tested binding of ZnF-UBPs to intact nucleosomes. ZnF-UBP domains of both USP16 and, surprisingly, USP49 bound to nucleosomes independently of ubiquitination whereas other ZnF-UBPs did not (**Fig. 3d, Extended Data Fig. 3e, f**). Indeed, USP49 displayed a 10-fold tighter interaction with unubiquitinated nucleosomes compared to USP16, with small differences in response units between ubiquitinated and apo nucleosomes possibly reflect secondary interactions with Ub (**Extended Data Figs. 2f, 3e, f**). We had not observed USP49 ZnF-UBP binding to UbCt or histone H4 tail peptides (**Fig. 1e, 3c**), suggesting the nucleosome interactions do not involve the UbCt pocket. We modelled the interaction between USP49 ZnF-UBP and an unubiquitinated nucleosome by AlphaFold3 as in a recent report^71^. In these models, USP49 ZnF-UBP interacts with histone H2B across (with some variation in orientation) (**Fig. 3e**), intriguingly at or near the H2B ubiquitination site Lys120, which is also the site deubiquitinated by USP49^32^. Consistent with the binding data, USP16 ZnF-UBP showed only weak nucleosome engagement, and AlphaFold3-modelling did not predict a consistent binding site. Moreover, the histone H4 tail is expected to be inaccessible in intact, DNA-bound nucleosomes, including the recombinant ones used in this study (see **Discussion**). AlphaFold3 models of full-length USP16 aligned to the recently determined structure^35^ (pdb-id 8wg5) indicate that the linker between catalytic and ZnF-UBP domains is sufficiently flexible to reach the histone H4 tail, if available (**Fig. 3f**).

### USP16 as a deFubiylase

The traditional view of the ZnF-UBP domain acting as a substrate binding domain, was further challenged when considering Fubi. The binding affinity to a Fubi-derived peptide (K_D_ = 59 µM) sat in between the histone H4 tail and Ub (**Fig. 3a**, **Extended Data. 3a**). USP16 is one of only two DUBs known to cleave Fubi^39^, and Fubi is not known to have roles as a stand-alone small protein, but rather is fused or conjugated. During Fubi biogenesis, it needs to be released from the ribosomal subunit eS30 (**Fig. 3g**). Poly-Fubi chains, or ubiquitinated Fubi, facilitated by the one Lys residue analogous to Lys27 on Ub^72^, have not yet been described. While Fubiylation has been observed on several substrates^39^, de-Fubiylation has also not been demonstrated. Therefore, unlike the histone H4 tail, the GlyGly motif in known Fubi substrates is never in a ‘free’ unattached state. Akin to a ubiquitinated substrate, the USP16 ZnF-UBP cannot mediate recognition of the Fubi C-terminus pre-cleavage (**Fig. 3g**). For both Ub and Fubi, this situation changes immediately post-cleavage, when Ub or Fubi C-termini are newly generated.

### Why is Ub the best substrate for the ZnF-UBP domain?

With a K_D_ of ∼19 µM, Ub remains the highest affinity substrate for the ZnF-UBP domain of USP16, which was a conundrum as it seemed unlikely that multiple DUBs act like USP5, by recognising and cleaving free Ub chains. The ascribed roles of USP16 as a histone DUB, together with the cellular concentration of ‘free’ Ub ranging between 10-70 µM^62,63^, were suggestive of regulatory mechanisms and ability of USP16 to respond to cellular Ub levels. We hence aimed to understand whether and how the ZnF-UBP domain affected the enzymatic activity of USP16 towards Ub as a substrate.

### The ZnF-UBP domain affects USP16’s kinetics and Ub binding profiles

We characterised and compared Ub binding and kinetic parameters of the USP16 catalytic domain (USP16 CD) with full-length USP16, and a full-length USP16 variant in which an R84A mutation in the ZnF-UBP domain abrogates Ub binding to the ZnF-UBP domain (**Fig. 4**, **Extended Data Fig. 4a-c**). Kinetic studies were performed using Ub rhodamine (Ub-Rhod), in which a rhodamine group fluoresces when enzymatically cleaved from the Ub C-terminus. Wild-type full-length USP16 showed the highest activity (K_M_ = 3.76 μM; k_cat_ = 0.39 s⁻¹). Strikingly, mutating or removing the ZnF-UBP domain resulted in enzymes with very similar, but much slower, kinetics. USP16^R84A^ and USP16 CD saturated at lower substrate concentrations with an increased apparent substrate affinity (K_M_ = 0.5-0.7 µM) but reduced kinetic turnover (k_cat_ = 0.05-0.06 s^−1^) (**Fig. 4b**). Overall, catalytic efficiency (k_cat_/K_M_) was similar across variants since k_cat_ and K_M_ changed proportionally. The Ub-Rhod substrate C-terminal Gly76 is obscured by the fluorescent extension and thus cannot bind the ZnF-UBP domain in full-length USP16 wild-type or USP16^R84A^ mutants; therefore, the difference in substrate affinity is not an effect of substrate binding. Instead, we hypothesised an effect on substrate release.

**Figure 4.**
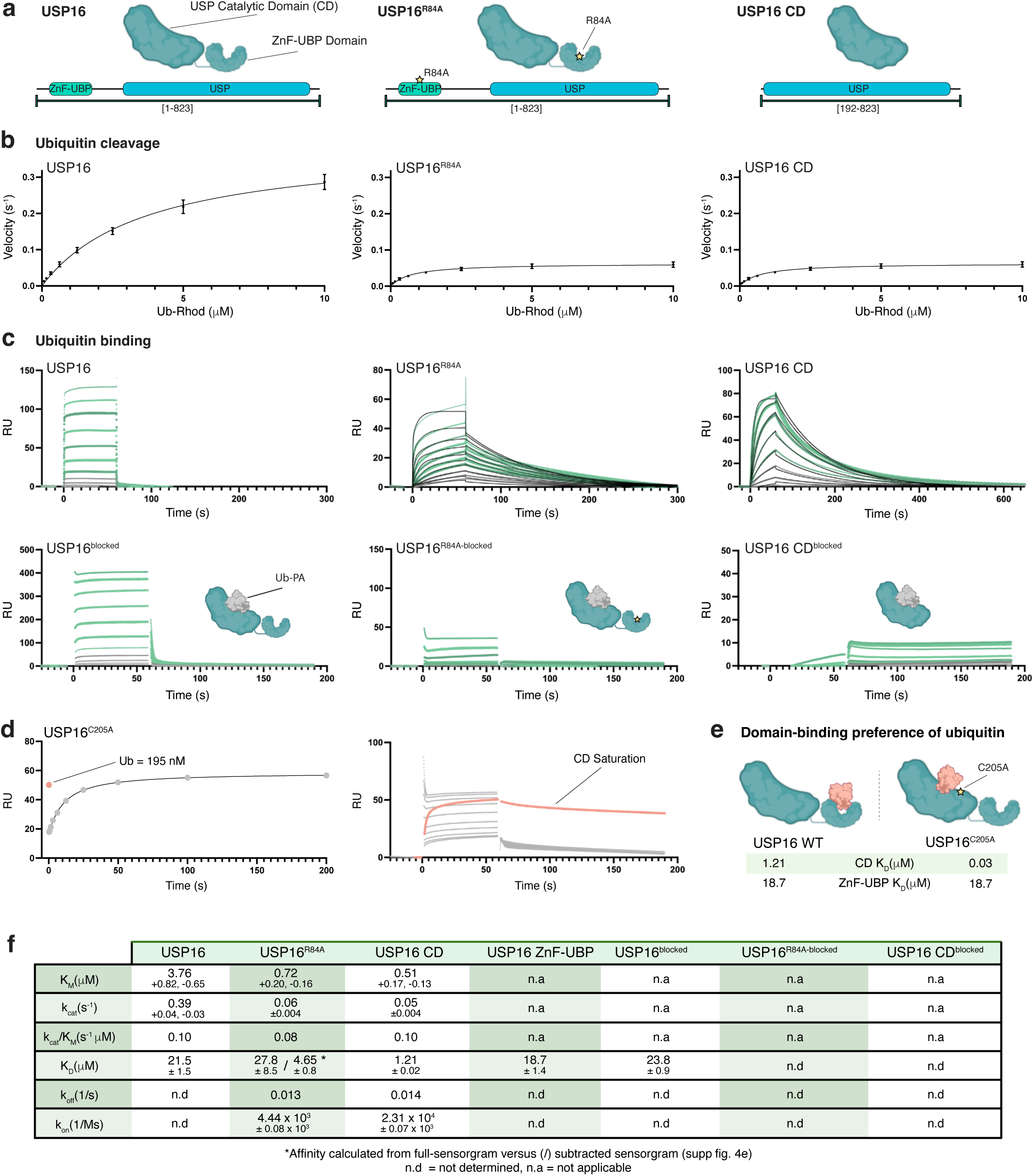
The ZnF-UBP domain of USP16 regulates enzyme catalysis. **a,** USP16 variants used in this study with construct boundaries shown in square brackets. See **Extended Data Fig. 4a-c** for quality controls. **b,** Michaelis–Menten kinetics of Ub-Rhod cleavage for USP16 variants. Plotted values represent mean and errors are standard error from the mean (s.e.m.) from *n =* 3 experiments. **c,** Ub binding sensorgrams for USP16 variants. ‘Blocked’ species were covalently modified with Ub-PA, which occupies the enzymatic S1 site (also see **Extended Data Fig. 4c**). Representative data shown from *n =* 3 experiments. Affinities were derived in **Extended Data Fig. 4d**. Kinetic fits used a 1:1 binding model. **d,** Binding of Ub to USP16^C205A^ where Ub binding and saturation of the catalytic domain is coloured orange. Increasing concentrations of Ub subsequently bind to the ZnF-UBP domain (in grey) Representative SPR data from *n =* 3 experiments. **e.** Ub binding preference to ZnF-UBP vs CD domain. **f,** Comprehensive activity and binding parameters for USP16 variants. Errors from enzyme kinetics represent s.e.m, errors from SPR data represent s.d. from the mean. All experiments were performed in triplicate.

We used SPR to study Ub interactions with USP16 and characterise product complexes of the enzymatic reaction. ‘Blocked’ enzymes were generated by reacting USP16 with irreversible Ub-propargylamide (Ub-PA) probes that covalently modify the active site Cys (Cys205 in USP16) in USP catalytic domains^73^ (**Extended Data Fig. 4c**). We found that USP16 interacted with Ub with a K_D_ of 21.5 μM, which was similar to the enzyme in the blocked state (K_D_ = 23.8 μM), and the isolated ZnF-UBP domain (K_D_ = 18.7 μM, see above) (**Fig. 4c, Extended Data Fig. 4d**). In contrast, USP16 CD interacted with Ub with 10-fold higher affinity (K_D_ = 1.21 μM) and showed no appreciable Ub binding in the blocked state (**Fig. 4c, Extended Data Fig. 4d, e**). Hence, despite the higher 1.21 μM affinity of the catalytic domain, the ZnF-UBP appears to dominate Ub recognition in the full-length enzyme. Intriguingly, USP16^R84A^ sensorgrams resembled a ‘hybrid’ of USP16 and USP16 CD, with unusual affinity parameters (**Fig. 4c, f**). Removing the highest concentrations from the USP16^R84A^ analysis yielded a similar curve to USP16 CD and increase in affinity (**Extended Data Fig. 4e**), suggesting some effects remain from the binding-incompetent ZnF-UBP in the full-length context. We were unable to observe similar effects with lower Ub concentrations in wild-type USP16, which showed rapid on- and off-rates, too fast for kinetic analysis (**Fig. 4c**). Strikingly, Ub binding for both USP16^R84A^ and USP16 CD showed significantly decreased dissociation rates (k_off_ = 0.013 and 0.014 s⁻¹, respectively) (**Fig. 4c**). These observations explain the cleavage kinetics, where ZnF-UBP ablated-species appeared saturated by Ub substrate, with accordingly low K_M_ values. We next attempted to disrupt the apparent dominance of the ZnF-UBP domain in determining enzyme kinetics. A catalytic C205A mutation increase USP affinity for Ub^74^, in USP16 ∼100-fold from 1.41 µM to 0.024 µM (**Extended Data Fig. 4f, g**). Ub binding in full-length USP16^C205A^ was now preferential to its catalytic domain which was immediately saturated by the first/lowest concentration of Ub in the SPR assay, and preceded Ub binding to the ZnF-UBP (**Fig. 4d**).

Collectively, our data reveal that the ZnF-UBP domain in USP16 is, surprisingly, a crucial regulator of enzyme activity, finely tuning Ub affinities in an enzyme with two independent Ub binding domains (**Fig. 4e**). Removing or mutating the ZnF-UBP domain, slows catalysis by creating tight Ub binding within the USP catalytic domain. In the intact enzyme the ZnF-UBP dominates Ub capture which enhances the catalytic rate (**Fig. 4f**).

### The ZnF-UBP domain affinity is finely tuned to the catalytic function of the CD

Based on our structural insights (**Fig. 1, 2**) we next mutated residues that would modulate ZnF-UBP binding affinity to Ub, to fractionally decrease but potentially also increase Ub binding (**Extended Data Fig. 5a**). The resulting panel of USP16 ZnF-UBP domains varied substantially in affinity relative to the wild-type domain, along predictions (**Fig. 5a, Extended Data Fig. 5b, c**). As in USP5 ^12^, a USP16 ZnF-UBP^D120A^ reduced Ub binding 3-fold (K_D_ = 76.1 µM). Intriguingly, increasing the hydrophobicity of the binding site, via E122Y or N85Y mutations, increased Ub affinity 5-and 10-fold, respectively, and combining both mutations was additive, generating a very high affinity ZnF-UBP domain (K_D_ = 0.7 µM). Combining N85Y and D120A mutations generated a domain with affinity in between the high affinity binders and wild-type (K_D_ = 7.7 µM) (**Fig. 5a**).

**Figure 5:**
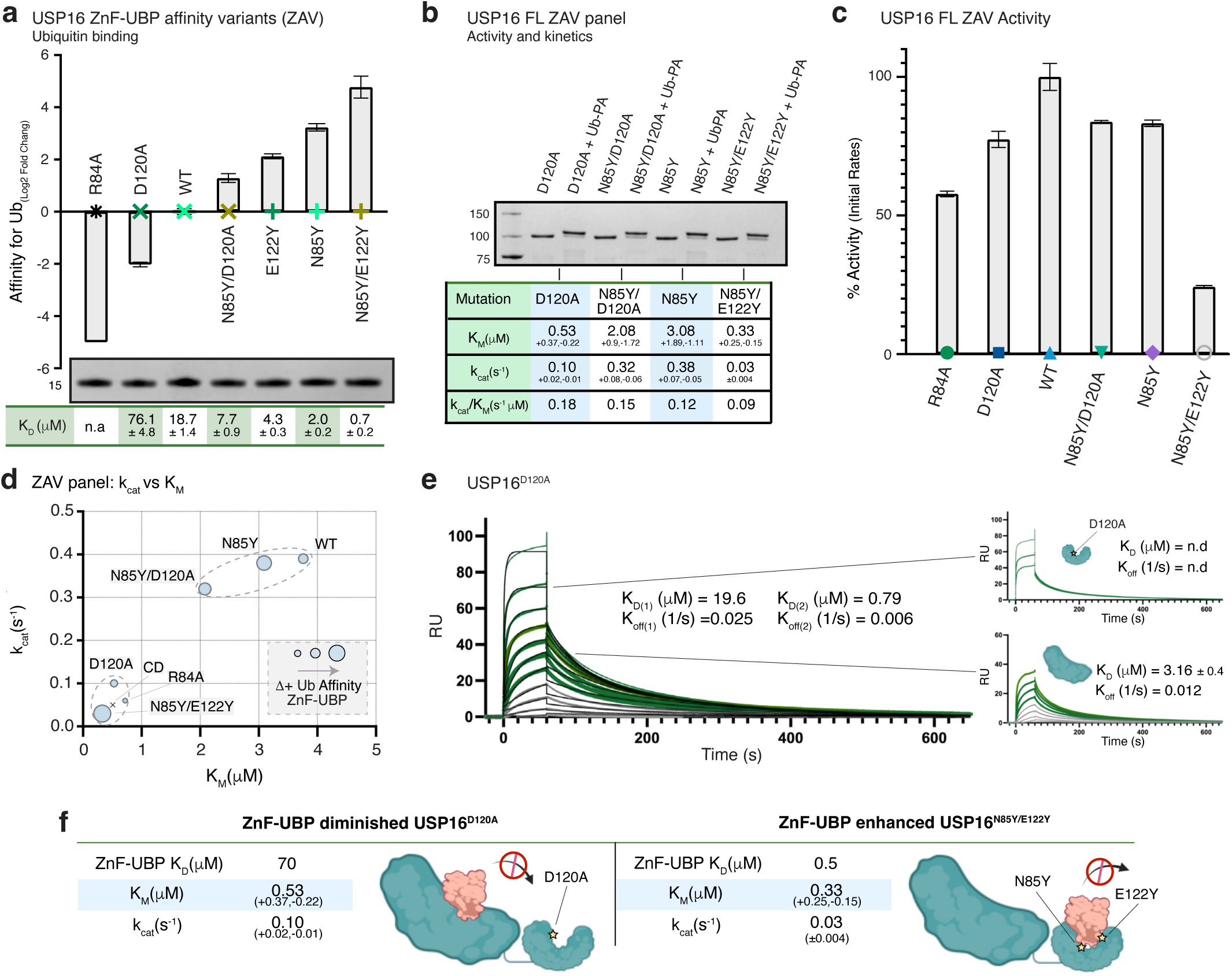
USP16’s ZnF-UBP domains affinity is finely tuned for kinetic activity. **a,** The ZnF-UBP affinity-variant (ZAV) panel, listing ZnF-UBP mutations, Ub affinity variation (*top*) and protein quality (Coomassie stained SDS PAGE gel, *bottom*). Affinities were calculated from curves in **Extended Data Fig. 5b** which correspond to the symbol colours shown in the main panel, and errors correspond to s.d. from the mean. **b,** Coomassie-stained Ub-PA modification assay for full-length USP16 ZAV panel. Kinetic parameters are tabulated, as determined from raw data in **Extended Data Fig. 5d. c,** Comparative initial rate % activity of each USP16 ZAV, which correspond to the symbol colours show in the summarised from raw curves in **Extended Data Fig. 5e**. Errors correspond to s.d. from the mean. **d,** K_M_ vs k_cat_ for all USP16 ZAVs grouped based on their ZnF-UBP affinity mutations relative to their activity. **e**, Ub binding sensorgram of USP16^D120A^ (*left*) fit with heterogenous ligand model. Zoom-in of the sensorgram (*right*) split into higher (*top*) versus lower (*bottom*) concentrations of Ub, showing distinct profiles for the ZnF-UBP^D120A^ domain and the CD, respectively. Kinetic parameters for separated catalytic domain profiles were determined in **Extended Data Fig. 5f**. A representative sensorgram from *n =* 3 experiments is shown. **f,** Comparison of the effects of low- and high-affinity ZnF-UBP domains on USP16 activity and binding.

Full-length variants with these mutations in the ZnF-UBP showed no difference in Ub-PA probe modification, yet showed changes in enzyme kinetics in Ub-Rhod cleavage assays (**Fig. 5b, Extended Data Fig. 5d**). Surprisingly, modulating ZnF-UBP affinity in either direction reduced enzyme activity compared to wild-type, showing how meticulously optimised the wild-type enzyme is (**Fig. 5c, Extended Data Fig. 5e**), while plotting K_M_ versus k_cat_ (**Fig. 5d**) revealed clear groupings across variants: The slowest variants (USP16^D120A^ and USP16^N85Y/E122Y^) have a markedly reduced catalytic rate and showed a pronounced shift in apparent substrate affinity, which we hypothesise drives the overall reduction in catalytic rate. In contrast, variants with smaller affinity increases (USP16^N85Y^ and USP16^N85Y/D120Y^) showed a modest yet real increase in substrate affinity and only a mild decrease in catalytic rate (**Fig. 5d**). Strikingly, Ub binding curves for the ZnF-UBP affinity-diminished species USP16^D120A^ revealed a binding curve most similar to the USP16 CD, with the sensorgram profiles of both domains visible (**Fig. 5e, Extended Data Fig. 5c, f**).

Hence, altering ZnF-UBP affinity for Ub appears to imbalance two finely-tuned domains, albeit via different mechanisms. A ZnF-UBP module with low affinity is unable to efficiently assist Ub release from the catalytic domain after cleavage, while ZnF-UBP domains with higher affinity hold on to Ub too, tightly blocking their ability to release newly-cleaved Ub from the catalytic site (**Fig. 5f**). Indeed, a unique ‘slow release’ sensorgram for USP16 ZnF-UBP^N85Y/E122Y^ supports this notion (**Extended Data Fig. 5b**). In either case, enzymatic substrate affinity is increased significantly, yet product release is hampered, creating a less efficient enzyme.

### Domain targeted Ub variants reveal distinct coordination between USP16 domains

A further question and complication in studying the USP16 enzymatic mechanisms concerns the control of free Ub concentration in the reaction; the generation of free Ub from enzymatic measurements can regulate each domain *in vitro*. To understand how free Ub might affect USP16’s activity, we performed Ub-Rhod assays in presence of excess free monoUb at 2 µM. Interestingly, adding monoUb led to product inhibition, which was most pronounced in USP16 wild-type (**Fig. 6a**). Notably, this is a scenario that clearly distinguishes USP16 and USP5 – in the latter, incorporation of free Ub accelerated the enzyme^12^. To disentangle domain-specific product inhibition, we engineered domain-selective Ub variants (**Extended Data Fig. 6, see Supplementary Text**) which were validated by SPR against both domains individually (**Fig. 6b**). Repeating the Ub-Rhod supplementation assay with these variants showed that most product inhibition occurred via the catalytic domain, with only a small contribution from the ZnF-UBP when using Ub^R3^ (**Fig. 6c, Extended Data Fig. 6e**). An initial 20 μM Ub^R3^ produced negligible effects, so we increased Ub^R3^ concentrations in the assay to 500 μM to ensure domain saturation yet the effect remained unexpectedly modest (**Fig. 6c, Extended Data Fig. 6e**). In contrast, Ub^G76V^ abolished activity, consistent with Ub becoming trapped in the catalytic domains and unable to be released by the ZnF-UBP. Mild effects were even seen with Ub^Dud^, likely due to the high concentrations used. Kinetic assays supplemented with Ub^R3^ showed an expected yet surprisingly mild decreases in K_M_ and k_cat_, with the impact of Ub^R3^ confined primarily to the initial the linear phase used for kinetic analysis. (**Fig. 6d**). The insignificant effects of Ub^R3^ were intriguing as they contrasted the earlier results that changes or deletion of the ZnF-UBP had pronounced inhibitory effects (**Fig. 4, 5**).

**Figure 6.**
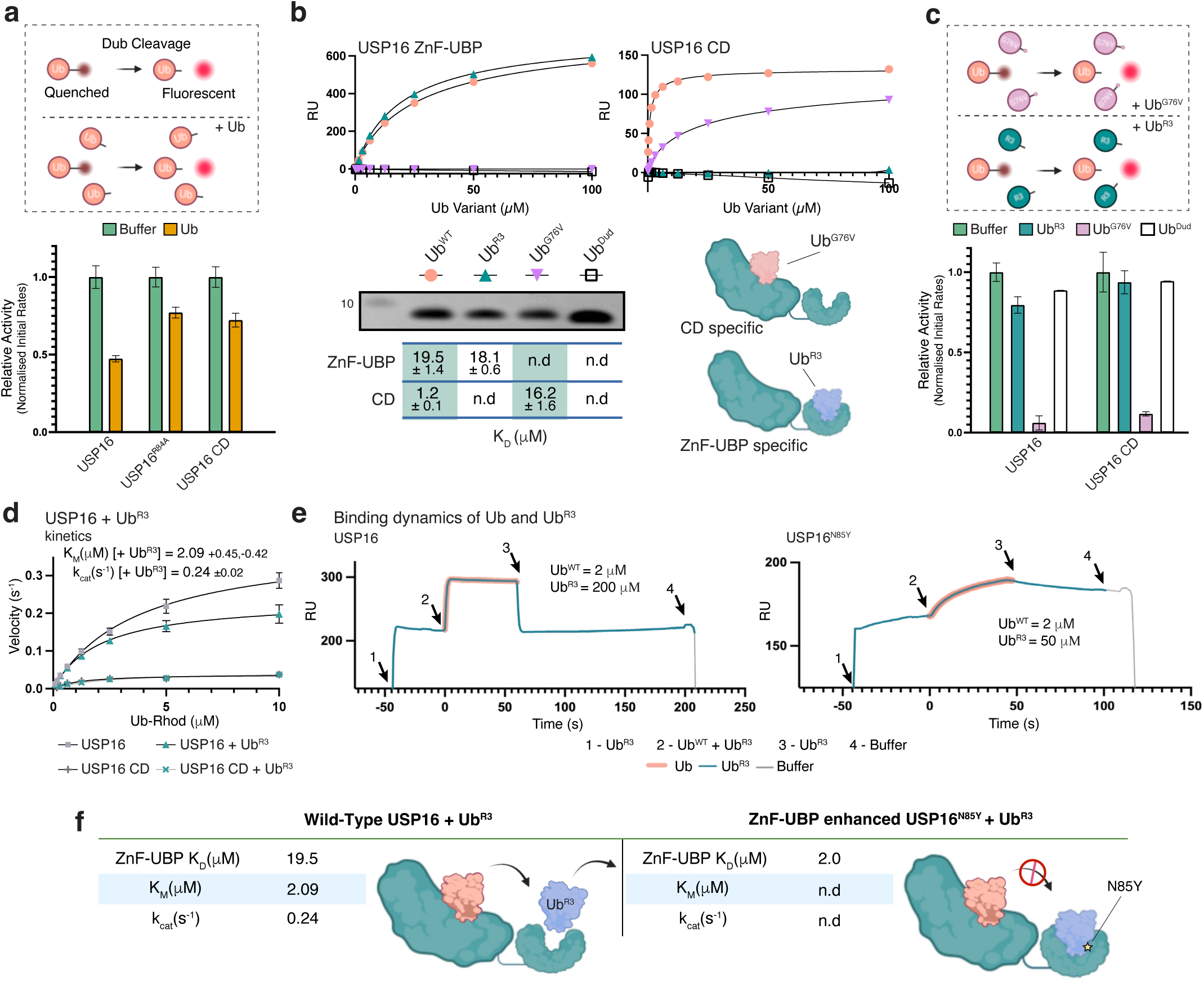
Engineered Ub variants reveal complex relationship between domains. **a,** Assay schematic and relative activity of Ub-Rhod (1 µM) cleavage by USP16 variants without (green, 100% activity) or in presence of 2 µM Ub (orange). Mean values are plotted, and error bars represent standard error from the mean from *n =* 3 experiments. **b,** Generation of domain specific Ub variants by rational design (see **Extended Data Fig. 6**), showing final set of Ub mutants, with binding affinities, and Coomassie stain. Affinities were determined by SPR (data from *n =* 3 experiments with representative curves shown). **c,** Relative activity of initial rates for Ub-Rhod cleavage by USP16 variants, shown as relative activity in presence of 500 µM Ub^R3^, Ub^G76V^, or Ub^Dud^ (from raw data in **Extended Data Fig. 6e**). Errors correspond to s.d. from the mean. **d,** Michaelis–Menten kinetics of Ub-Rhod cleavage for USP16 variants supplemented with Ub^R3^ at 20 µM. Plotted mean ± s.e.m. from *n =* 3 independent assays. **e,** SPR ABA injection using combinations of Ub and Ub^R3^ binding to USP16 and USP16^N85Y^. USP16 and variants were attached to the SPR surface via a biotinylated Avitag (see **Methods**). A representative experiment is shown, from *n =* 3. **f,** Summary of the effects of Ub^R3^ supplementation on USP16 activity.

We hence decided to test binding dynamics between both domains. Ub^R3^ did not measurably affect Ub binding to the catalytic domain when supplemented alongside a standard Ub binding assay, and also retained binding to the ZnF-UBP of full-length USP16 (**Extended Data Fig. 6f**). We therefore designed an ABA-format SPR injection to observe any synergistic or antagonistic effects between both domains (**Fig. 6e**). Briefly, the system allows injection of two analytes (analyte mix A and analyte mix B) to sequentially study complex systems. Saturating USP16 with Ub^R3^ (*Injection A*) was expected to block the ZnF-UBP. This was followed by *Injection B* in which identical Ub^R3^ was mixed with Ub^WT^, which we expected to reveal CD binding curves (as in **Fig. 4c**). However, the resulting sensorgram was purely additive, with all signal attributable to ZnF-UBP binding from both Ub^WT^ and Ub^R3^ (**Fig. 6e, Extended Data Fig. 6h**), with the ZnF-UBP domain fast releasing Ub^R3^ to displace Ub^WT^ from the CD.

We hence used the tight binding and slower-off rate USP16^N85Y^ mutant (**Extended Data Fig. 6g**), which retained Ub^R3^ in the ZnF-UBP domain. When we repeated the ABA experiment with USP16^N85Y^, which finally revealed binding parameter contributions of the CD upon Ub^WT^ addition, with the expected slow release of Ub^WT^ from the catalytic domain observable as the mutated ZnF-UBP^N85Y^ remained saturated with Ub^R3^ (**Fig. 6e**). Higher Ub^R3^ concentrations were required to uncover catalytic domain contributions to Ub^WT^ binding in the USP16^N85Y^ mutant (50 µM Ub^R3^ vs. 2 µM Ub^WT^), and we hence do not expect Ub to out-compete Ub^R3^ at the ZnF-UBP. Therefore, when the ZnF-UBP is occupied by Ub^R3^, wild-type Ub binds in the catalytic domain and is released with rather slow kinetics without ZnF contributions (**Fig 6f**).

The fast-binding kinetics in the wild-type enzyme enable USP16 to remain vigilant about whether product needs to be released. Reducing the off-rate in the ZnF-UBP domain also reduces off-rate, i.e. product release, and hence kinetics, of the catalytic domain. This model derived for USP16 differs considerably from previous ideas about the role of the ZnF-UBP domain in USP5 as a substrate anchor^13^, or from a model whereby a change in activity is simply due to steric hindrance of the catalytic domain due to a ZnF-UBP-bound second Ub.

### Overcoming product inhibition selectively increases DUB activity

The observations using Ub variants to control domain occupancy hinted at important roles for product inhibition and Ub concentrations in kinetic assays. We considered an alternative way to overcome product inhibition in USP16. Here, to remove all free Ub as it is produced by the enzyme, we included the high-affinity, tight binding USP16 ZnF-UBP^N85Y^ as a sequester domain (SD) in Ub-Rhod assays (**Fig. 7a**). The SD does not interact with Ub-Rhod, and hence the assay substrate has only one binding site in the enzyme, namely the catalytic domain. In contrast to Ub product inhibition (**Fig. 6a, reproduced** for comparison in **Fig. 7a**), inclusion of SD increased the catalytic rate across all USP16 variants, with a more pronounced effect in ZnF-UBP-ablated variants (**Fig. 7a**). It was interesting that incorporation of exogenous SD doubled the activity of the catalytic domain, likely by helping to release, or prevent re-binding of, post-cleavage Ub products from the active site.

**Fig. 7.**
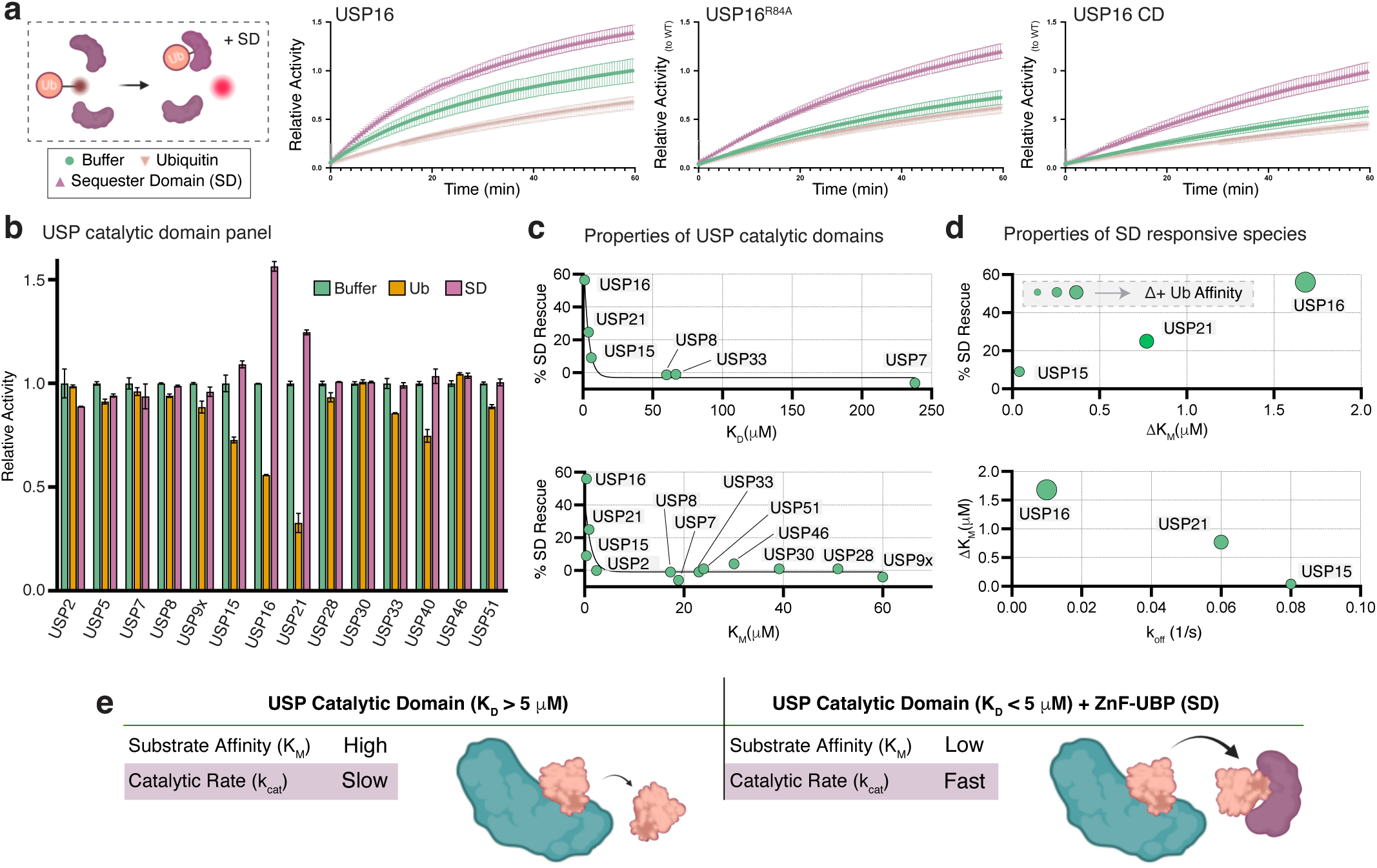
USP activity can improve with addition of a ZnF-UBP ‘Sequester Domain’ in *trans*. **a,** Assay schematic, indicating the use of the high-affinity USP16 ZnF-UBP^N85Y^ ‘sequester domain’ (SD) as an assay additive (purple). Activity curves for USP16 variants (see **Fig. 4**) in presence of 2 µM Ub or 2 µM SD. Mean values are plotted, and error bars represent standard error from the mean from *n =* 3 experiments. **b,** Relative activity of Ub-Rhod cleavage by panel of USP catalytic domain constructs (see **Supplementary Table 1**) in presence of 10 or 2 µM Ub or 2 µM SD. Raw data are shown in **Extended Data Fig. 7a, b**. Error bars correspond to standard error of the mean from n=3 experiments. **c,** *Top:* Percentage increase in activity following SD supplementation versus measured affinities of USP domains for Ub (see **Extended Data Fig. 7a-c**). *Bottom:* Percentage increase in activity following SD supplementation versus both measured and reported K_M_ values. **d,** *Top:* Percentage increase in activity following SD supplementation against ΔK_M_ for enzymes which were improved by addition of SD. *Bottom:* ΔK_M_ against k_off_ for enzymes which were improved by addition of SD. See **Extended Data Fig 7. e, Summary** of scenarios in which SD can improve activity of USP catalytic domains.

We next wondered whether the observed activity increase would be a global feature in USP enzymes, and repeated the Ub/SD supplementation assay across a panel of USP domains (**Fig. 7b, Extended Data Fig. 7a**,b). Notably, Ub did not inhibit—and SD did not rescue—as broadly as anticipated; only three USP catalytic domains, USP16, USP15 and USP21, showed increased activity after incorporation of SD (**Fig 7b, Extended Data Fig. 7a**). We measured Ub-binding affinity across the panel and observed a correlation: enzymes that were rescued by the ZnF-UBP tended to have high Ub affinity (where a K_D_ curve could be determined), while monoUb binding was not observable in catalytic domains of USP2, USP5, and USP9X (**Fig. 7c**, *top***; Extended Data Fig. 7a-c**). This trend also tracked with K_M_ for species in which it could be determined/which values were available in the literature^38,75–78^ (**Fig. 7c**, *bottom***, Extended Data Fig. 7a, d, Supplementary Table 1**). Among species with measurable effects, ΔK_M_ exhibited an approximately linear relationship across the affected enzymes proportional to % SD rescue and increasing affinity for Ub (**Fig. 7d**, *top*), while a similar inverse trend was observed for off-rates (**Fig. 7d**, *bottom*), although off-rates could only be effectively determined for the SD-affected species, leaving the correlation uncertain beyond these.

Our data suggest that the SD, a high affinity Ub binding domain able to soak up all free Ub product in the assay, lowers the apparent USP domain K_M_ for Ub; this effect is observed only in enzymes whose catalytic domains bind Ub with high affinity and/or have slow off rates (**Fig. 7e**).

### ZnF-UBP domain positioning enables intra-molecular contributions to the catalytic mechanism

A kinetic model exploiting connected domains was corroborated by USP16 AlphaFold3 prediction that shows an interface between the ZnF-UBP and catalytic domain which places an S1-site bound Ub (or, Fubi, as in **Fig. 3g**) tail into close proximity to the ZnF-UBP UbCt binding pocket (**Fig. 8a**). In silico Ala scanning identified residues that stabilise inter-domain interactions (**Fig. 8a, Extended Data Fig. 8a**). “Binding-interface” (BI) mutants, USP16^Y295A/S756A^, in full-length and CD variants were active (**Extended Data Fig. 8b**). Indeed, the mutation in the full-length enzyme reduced activity suggesting domain communication was affected (**Fig. 8b**). Surprisingly, BI mutation also affected USP16 CD, which, strikingly, showed improved enzyme kinetics in Ub-Rhod assays compared to wild-type USP16 CD (**Fig. 8b, Extended Data Fig 8c-e**). Thermostability profiles showed no defects (**Extended Data Fig. 8f**) and results were confirmed with fresh preparations. It appears that the relative proximity of Tyr295 to the catalytic site may affect Ub binding kinetics; as we had shown, Ub off-rates are rate-limiting. Indeed, SPR Ub binding studies showed a slight but consistent decrease in affinity and off-rates for the catalytic domain BI mutant (**Fig. 8c**, **Extended Data Fig. 8g**).

**Fig 8.**
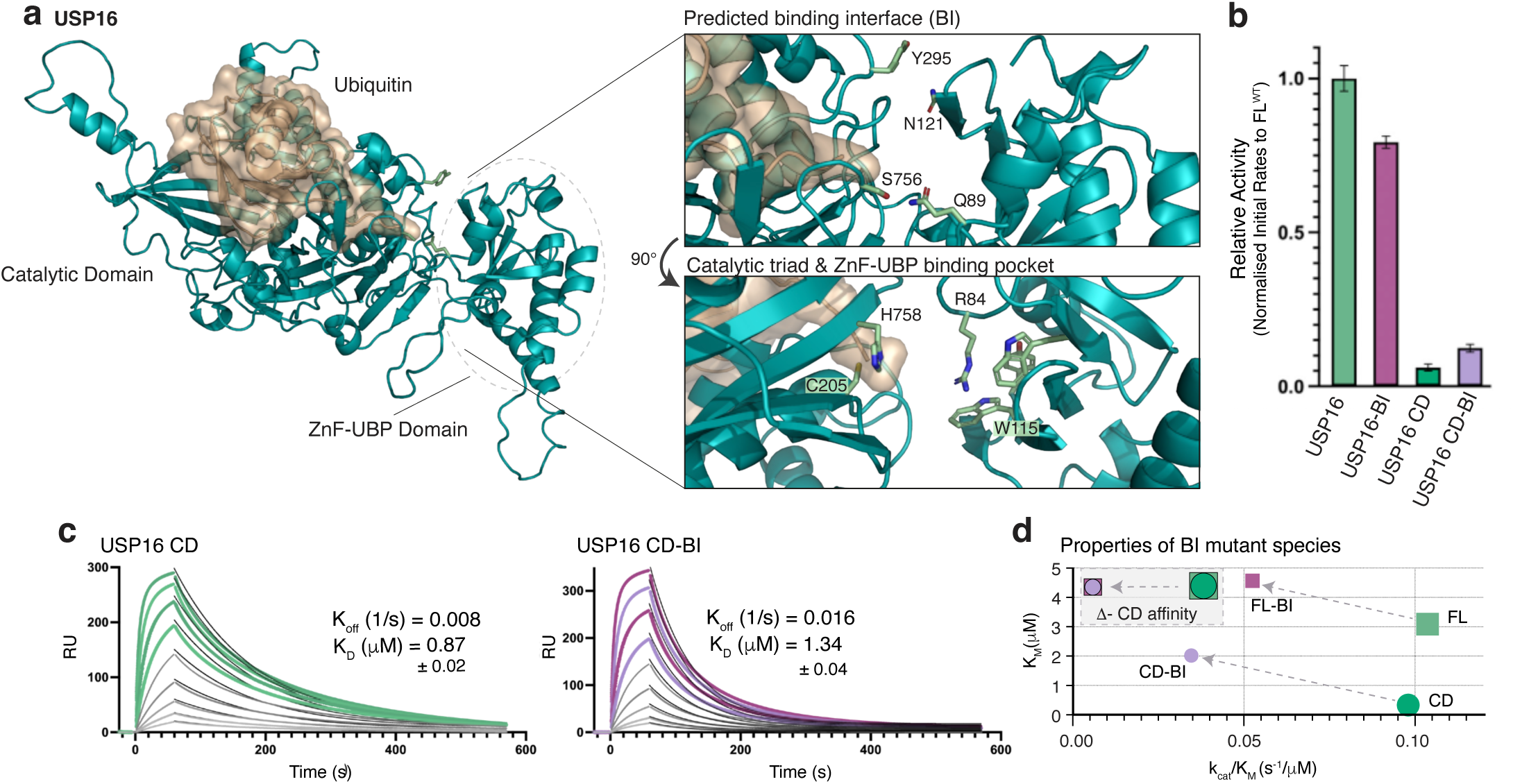
A binding interface between the ZnF-UBP and catalytic domain allows them to process Ub enables domain communication. **a,** AlphaFold3-predicted model of full-length USP16 (*left*). The disordered insertion (aa 397 – 627) was placed sporadically^37^ and has been omitted for clarity. Close-ups (*right*) show the predicted binding interface between the ZnF-UBP and CD (*top*), and the proximity of the catalytic triad to the ZnF-UBP binding pocket (*bottom*). **b.** Relative activity of Y295A/S756A binding interface (BI) mutants derived from raw data in **Extended Data Fig. 7c**. Error bars correspond to standard error of the mean from n=3 experiments. **c.** Sensorgrams for USP16 CD WT and BI variants fit using a 1:1 dissociation model. Representative sensorgrams shown from *n*=3 experiments. Errors represent s.d. from the mean. **d.** K_M_ vs. catalytic efficiency (k_cat_/K_M_) for USP16 BI panel grouped by Ub binding affinity.

This last insight suggested yet another potential mechanism to regulate USP domains. Tyr295 in USP16, is a well conserved bulky hydrophobic residue within ‘Box 2’ of USP catalytic domains^7^, just upstream of the Gln-rich region that interacts with the Ub tail (**Extended Data Fig. 8h**). A hydrophobic residue in this position may be important in substrate binding, and also interacts with nucleosomes (**Extended Data Fig. 8i**). Plotting catalytic efficiency (k_cat_/K_M_) against substrate affinity (K_M_) highlighted a consistent decrease in apparent substrate affinity for both full-length and USP16 CD variants, accompanied by a reduction in catalytic efficiency, a behaviour not seen with previous ZnF-UBP affinity mutants (**Fig. 8d, Extended Data Fig. 8e**; compare **Fig. 5b, d**). Together, this data suggests that the predicted intra-domain interface communicates and regulates catalytic efficiency; while this could be an intrinsic effect of catalytic domain stability indirectly promoting product release, the lower activity could also originate from impaired domain communication, preventing the ZnF-UBP domain to perform efficient product release.

## Discussion

We here provide a comprehensive characterisation of a common, yet specialised, UBD which features as an exo-domain in ∼20% of human USP DUBs. To date, exo-domains are mostly known for their ability to provide substrate specificity to non-specific USP domains. A 20-year old study on USP5^12^ revealed its ZnF-UBP domain to bind the unattached C-terminus of Ub, which catalytically activated the enzyme to cleave free Ub chains into monoUb. However, it was unclear whether this mechanism applied to other DUBs. It was also an attractive model that ZnF-UBP domains could ‘read’ the cellular Ub concentration, and become active once Ub pools are depleted, to release Ub from histones. Our comprehensive and comparative biochemical and structural annotation of human ZnF-UBP domains, while reconciling much of the fragmented literature, propose an alternative role for ZnF-UBP domains, as delineated for the nucleosome DUB USP16.

### ZnF-UBP domains in substrate binding

C-terminal GlyGly motifs are not only found in Ub and other UBLs, but intriguingly, also other proteins such as histone H4. Indeed, USP16 binds to histone H4 tail, albeit with high µM affinity; this tail is not available in intact nucleosomes. Perhaps, USP16 targets open, unwound or partially assembled nucleosomes^79^, which must exist as modifications of the histone H4 tail have been reported^80,81^. These ideas need structural validation by nucleosome specialists. However, the role of the ZnF-UBP domains in substrate interactions may not be generally true. The more important binders of ZnF-UBP domains could be those generated by the catalytic domain, the Ub (and Fubi) molecules in the S1 site of the enzyme.

### The ZnF-UBP domain as a crucial component of the catalytic cycle

Indeed, without a functional ZnF-UBP domain, USP16 catalysis was significantly reduced. Kinetic and binding experiments revealed that this effect is due to a catalytic domain that apparently has no efficient mechanism to release the cleaved product. Transferring this important product release task to an exo-domain is striking; we are not aware of other enzymes in the Ub system, or any other system, where this has been demonstrated. As such, our data highlight that a holistic characterisation of full-length DUBs is necessary. We note that many DUBs across families comprise UBD exodomains that may impact product release, differing significantly from long-held ideas in the field^15^. However, with the largest Ub interaction interface and highest affinity^5^, slow product release may be a special feature of USP enzymes. Our work also identifies residues in an additional allosteric switch that appears to contribute to slow enzymatic off-rates, and demonstrated that apparent DUB activity can be stimulated in trans, by supplying a product-capture domain in enzymatic reactions. The latter finding may have practical applications, and could perhaps even suggest the presence of exo-factors that ‘activate’ the enzyme by increasing product turnover.

### The complete catalytic cycle of USP16

We expect that the two roles of the ZnF-UBP domain, in substrate binding (e.g. through the histone H4 tail), and in product release, collaborate and are both important in the full catalytic cycle of USP16. In an apo enzyme, the ZnF-UBP domain would be available to bind substrates, including histones, albeit weakly. Once Ub is cleaved in the S1 site, the ZnF-UBP domain can binds its *primary* substrate, a freshly cleaved Ub C-terminal tail. Fast off-rates from the ZnF-UBP domain prevent that the enzyme being blocked at either step. As explained, such structural considerations are feasible and indeed supported by existing inter-domain allostery (**Fig. 8**). Indeed, the dynamic model proposed also suggests in-built ‘release’ mechanisms for USP16, where the ZnF-UBP domain, drawn to its primary GlyGly substrate, dislodges from the deubiquitinated nucleosome, and releases Ub in a swift single action.

### The biological context – a nucleosome (or a histone) DUB responding to Ub concentration?

Finally, our data may provide fresh insight into the role of USP16 in the cell. The well-established function of USP16 as a histone DUB was not least confirmed by the recent cryo-EM structure; although additional mechanisms of specificity must exist to target USP16 to select loci^27,43^. A further question is whether and how USP16 responds to changing Ub concentrations in the cell. We here show that USP16 (and some but not many other USP DUBs) are prone to product inhibition, even at low Ub concentrations easily observed in cells. The presence of ∼20 µM ‘free’ Ub is not typically considered in DUB assays performed in vitro. Exo-domains in DUBs could exacerbate, neutralize or inhibit interactions with free Ub, and could affect enzyme activity in context of the cellular Ub pool^5,82^. Such scenarios appear understudied in the current literature.

The Ub field has long been intrigued by mechanisms in which the concentration of free Ub is sensed^61^, with ZnF-UBP domains implicated from their first characterisation as free Ub binders^12^. In a broader sense, however, all USPs, all DUBs, and indeed any machinery that interacts with free Ub both shape, and are shaped by, the Ub pool itself. Enzyme kinetics and protein interactions are inherently concentration and affinity-dependent, making the Ub pool not just a background reservoir but an active, dynamic participant in ubiquitination and deubiquitination^55,61^. Thus, beyond the mechanistic detail of (a) specific free Ub sensor(s), we need to consider changes in the Ub pool when studying enzyme kinetics. In this way, Ub metabolism is perhaps best understood as a self-regulating, synergistic system in which the Ub pool and its regulators are inseparably intertwined.

## Acknowledgments

The authors would like to thank members of the Ubiquitin Signalling Division at WEHI for reagents, advice and comments on the manuscript, especially Simon Cobbold, Nicholas Kirk, Winnie Tan, Richard Birkinshaw, and Stephin Vervoort, as well as Shabih Shakeel and Peter Czabotar for their contributions to project ideation. We also acknowledge Cody Hall and Bernhard Lechtenberg for technical help and reagents, and Ha Na Kim (Cytiva) and Alexander Fish (Netherlands Cancer Institute) for their guidance in designing SPR experiments. This work was funded by WEHI and an NHMRC Investigator Grant (GNT1178122 to DK). JAA is supported by an Australian Government Research Training Program Scholarship.

Research was further supported by an NHMRC Independent Research Institutes Infrastructure Support Scheme grant (361646) and Victorian State Government Operational Infrastructure Support grant. This research was undertaken in part using the MX2 beamline at the Australian Synchrotron, part of ANSTO, and made use of the Australian Cancer Research Foundation (ACRF) detector.

## Author contribution

JAA designed, performed, and analysed all experiments in this manuscript, and wrote the manuscript with DK. RA designed, performed and analysed NMR experiments, PS performed activity assays for the DUB panel, AC purified numerous protein constructs, UN co-supervised the project at early stages, JJB provided advice and analysis on enzyme kinetics and SPR experiments, and DK contributed to design, analysis, and manuscript writing, and obtained funding.

## Online Methods

### Construct preparation

All ZnF-UBP and USP16 FL and CD variants were sequence-optimised for expression in *E. coli* and ordered as gBlocks Gene Fragments (IDT Singapore). gBlocks were inserted into pOPINK vectors^85^ with N-terminal GST tag and 3C cleavage site using In-Fusion cloning (Takara Bio) (see **Supplementary Tables 2, 3**). N-terminal AviTags™ were incorporated into specified variants, and several USP16 species were ordered in custom pET17b plasmids (containing the same tag system) through VectorBuilder. Mutations were introduced using the Q5 site-directed mutagenesis kit (NEB). All plasmids were grown and prepared in DH5α *E.coli* strain (NEB) using standard molecular biology techniques.

### Protein expression and purification

Constructs were expressed in BL21(DE3) *E. coli* (NEB) in an overnight starter culture supplemented with appropriate antibiotics and shaking at 200 rpm, which was maintained for all growth steps. Starter cultures were added 1:40 into 1 L of 2xYT media and grown at 37°C until an OD_600_ of between 0.5-0.8 was reached, followed by overnight induction with 0.5 mM IPTG at 20°C. All subsequent steps were performed at 4°C and/or on ice. Cultures were centrifuged at 5000 × g and pellets were either snap-frozen with liquid N_2_ or lysed by sonication in 20-50 mL lysis buffer (50 mM Tris, 5 mM DTT and varying concentrations of NaCl, glycerol, and pH, see **Supplementary Tables 2, 3**) supplemented with Complete protease inhibitor tablet (1 per/L), 0.1 mg/mL DNase, 1 mg/mL lysozyme, and 1 mM MgCl_2_. Cells were lysed by sonication (70% amplitude, 3 s on 6 s off) then cleared by centrifugation at 40,000 × g for 30 min and 0.45-μm filtered. Cleared lysate was incubated with 0.5–3 ml buffer-equilibrated Glutathione Sepharose 4B resin (GE Healthcare) for 1 h and washed with a 500 mM NaCl lysis buffer variant. ÄKTA pure systems (GE Healthcare) were used for subsequent purification steps. ZnF-UBP variants were purified by size exclusion chromatography (SEC) on a HiLoad 16/600 Superdex 75 pg (Cytiva) column in 20 mM HEPES, 1 mM TCEP with varying NaCl, glycerol and pH conditions (**Supplementary Tables 2, 3**).

Full-length USP16 and catalytic domain variants were lysed in 50 mM Tris, 500 mM NaCl, 1 mM DTT and 5% glycerol (v/v) at pH 8.0 as above. For biotinylated AviTag^TM^ constructs, 196 µM biotin, 1 mM DTT, 2.9 mM ATP and 50 µL recombinant His-BirA (40 µM) at (kindly supplied by Dr. Bernhard Lechtenberg and Anthony Cerra) were added during the overnight incubation. For ‘blocked’ variants containing a Ub-PA probe in the active site, 1 mg of Ub-PA (see below) was added during incubation. All variants were purified via SEC on a HiLoad 16/600 Superdex 200 pg (Cytiva) in 20 mM HEPES, 1 mM TCEP, 150 mM NaCl at pH 8.0. USP16 full-length variants were diluted to 50 mM NaCl and further purified on a MonoQ column over a 500 mM NaCl gradient, eluting at ≈ 165 mM NaCl. Ub and Ub variants were produced from pET17b plasmids as described previously^86^. For Ub-PA, Ub (1–75) was expressed in pTXB1 vectors and prepared as described elsewhere^87^. All purified proteins were concentrated at 4000 × g by Amicon centrifuge filtration (3 kDa cutoff for ZnF-UBP and Ub, 30 kDa for USP16 variants), flash-frozen in liquid N_2_ in aliquots, and stored at −80°C.

For NMR sample preparations, USP16 ZnF-UBP (aa 22–144) transformed cells were grown in M9 medium supplemented with ¹⁵NH₄Cl (Cambridge Isotopes Laboratories, Inc.). Cells were induced at an OD_600_ 1.0 with 1 mM IPTG for 4 h at 37 °C. Harvested cells were processed as normal excepting the SEC buffer contained 25 mM sodium phosphate, 100 mM potassium chloride, 2 mM DTT, pH 7.0 to match previous NMR conditions^22^. Purified protein was used immediately for the data collection.

### Quality control

Protein quality was assessed by SDS PAGE using 4-12% NuPAGE gradient gels (Invitrogen) at multiple stages during purification. In addition, samples were diluted to between 1-10 µM in 20 mM HEPES and 150 mM NaCl, pH 8.0, and thermostability was measured on the Tycho.NT6 (NanoTemper Technologies) using the default protocol. Data were collected and analysed using the Tycho.NT6 inbuilt analysis software. To confirm activity of catalytically competent USP16 variants, and to generate ‘blocked’ variants, we utilised Ub-PA covalent probes^73^. Here, 0.5 µM of DUB was incubated with 2 µM of Ub-PA either at room temperature (RT) or 37 °C and at specified time intervals.

### Crystallisation of USP16 and USP49 ZnF-UBP domains

Crystallisation screens of USP16 ZnF-UBP (aa 22–144; 15.6 mg/mL) and USP49 ZnF-UBP (aa 1–103; 23 mg/mL) were conducted at the CSIRO C3 Collaborative Crystallisation Centre in Melbourne, Australia. Shotgun and PACT screens were performed by sitting-drop vapour diffusion in 300 nL drops at two concentrations (maximum and half-maximum) and two temperatures (20 °C and 8 °C). Crystals appeared under multiple conditions. Thin, rod-like crystals of USP16 ZnF-UBP (30% (w/v) PEG 4000 at 20 °C) and USP49 ZnF-UBP (0.2 M sodium chloride; 2 M ammonium sulphate; 0.1 M sodium cacodylate (pH 6.5), 8 °C) were harvested, cryoprotected in mother-liquor reservoir solution supplemented with 20% (v/v) glycerol, and vitrified in liquid N_2_.

### Crystallographic data collection, phasing and refinement

Diffraction data was collected at the Australian Synchrotron (ANSTO) using the MX2 beamline (λ = 0.9537 Å, 100 K)^88^ and processed with XDSme (v.0.6.5.2)^89^. Data was merged in AIMLESS^90^ through CCP4i^91^. Phaser was used for molecular replacement^92^ within Phenix^93^ using pdb-id 2I50 for USP16 and 3C5K for USP49 as models. Coot^94^ was used to build the model with iterative rounds of refinement performed in Phenix. The USP16 ZnF-UBP structure contained three copies within the asymmetric unit, with insufficient density in chains A and B between aa 31 to 36. Following refinement, atomic coordinates from Chain C were transplanted to these regions and used in further model building and refinement. Some flexible regions (aa 55 – 69) were not modelled due to apparent disorder. The Ramachandran statistics for USP16 ZnF-UBP showed no outliers, 3.79% of residues in allowed and 96.21% in favoured regions. USP49 ZnF-UBP Ramachandran statistics contained 0.97% outliers, none allowed and 99.03% favoured. Structure figures were generated in PYMOL^95^.

### NMR measurements

NMR titration experiments were recorded at 298 K on a 600 MHz Bruker Avance III spectrometer equipped with a TCI cryoprobe. All samples were with 10 % D₂O in 5 mm Shigemi microtubes, as previously reported^22^. For titration, 135 µM ^15^N-labelled USP16 ZnF-UBP was titrated with H4 C-terminal peptide at 270, 405, and 675 µM concentrations, equivalent to 1:2, 1:3 and 1:5 ratio. Data were processed with TopSpin 4.1 and NMRPipe^96^ and analysed in POKY^97^ to quantify the chemical-shift perturbations (CSP) using the formula:

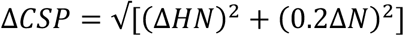

### Structural prediction

Structural models of specified ZnF-UBP and full-length USP16 variants were predicted using the AlphaFold3 webserver^98^. The predicted models with the highest LDDT scores are shown. PAE viewer was used to generate prediction quality maps^99^.

### Binding studies using fluorescence polarization (FP)

All peptides were purchased from Genscript^TM^ and were N-terminally FITC-labelled in the format of ‘FITC-Ahx-SEQUENCE’. Peptides were dissolved to 20 mM in either DMSO or Milli-Q water as advised by solubility reports. For assays, peptides were held at 5 nM, and ZnF-UBP domains titrated in a 1:2 dilution series up to the maximum achievable concentration, performed in FP assay buffer (10 mM HEPES, 100 mM NaCl, 0.5 mM TCEP, 0.5 mg/mL BSA, pH 6.0–8.0). Assays were performed in black, flat-bottomed 384-well plates (10 µL volume) and read on a CLARIOstar Plus (BMG Labtech) using default FP parameters (excitation α=482 ± 8 nm; dichroic LP α=504 nm; emission α=530 ± 20 nm). Data were analysed with MARS data analysis software.

### Binding studies using surface plasmon resonance (SPR)

All experiments were performed on either a Biacore 8K or Biacore S200 instrument (Cytiva). CM5 sensor chips (Cytiva) were used for amine coupling with the Amine Coupling Kit (Cytiva). Ligands were diluted to 50 µM (Ub and ZnF-UBP) or 10 µM (USP16 variants) in immobilization buffer (150 mM NaCl, 20 mM HEPES, pH 6.8–8.0), then further diluted 1:10 to a final volume of 120 µL in 50 mM sodium acetate (pH 4.5 for Ub, pH 5.0 for ZnF-UBPs, and pH 5.5 for USP16 variants). For the USP panel, ligands were diluted to the concentrations indicated in **Supplementary Table 1**. For streptavidin coupling, biotin-labelled Avi-tag ligands were diluted to 5 µg in immobilization buffer (150 mM NaCl, 20 mM HEPES, pH 6.8–8.0) and immobilised on an SA sensor chip (Cytiva). Immobilization reagents (500 mM NaCl, 50 mM NaOH; and 500 mM NaCl, 50 mM NaOH, 50% (v/v) isopropanol) were prepared according to the instrument method protocol. For all nucleosome binding experiments, biotin labelled nucleosomes (Epicypher) were immobilised as above. For kinetic measurements, immobilization levels for the R_ligand_ were adjusted to achieve an Rmax_Analyte_ < 250 RU.

Immobilised chips were primed with running buffer (150 mM NaCl, 20 mM HEPES, 1 mM TCEP, 0.005% (v/v) Tween 20, pH 6-8). Analytes were diluted in the same buffer and titrated at the desired concentrations (25 – 500 µM) into 96-well round-bottomed plates. Flow rates of 30 µL/s were used, with contact times for each species indicated in the corresponding figures. Affinity and kinetics analyses were performed using either Biacore S200 Evaluation Software or Biacore Insight Evaluation Software. For ABA injections, Ub/Ub^R3^ combinations were prepared with injection times indicated in the figures.

### Kinetics assays

Ubiquitin-rhodamine (Ub-Rhod, UbiQ) was dissolved at 1 mM in DMSO and diluted in kinetics assay buffer (20 mM Tris, 0.03% (w/v) BSA, 0.01% (v/v) Triton X-100, 0.03% (w/v) EDTA, 100 mM NaCl, pH 8.0) to prepare the final titration. Black, flat-bottomed 384-well plates were set up with USP16 or its variants at a constant final concentration of 4 nM. For kinetics calculations, a standard reaction containing 100 nM USP16 and 1 µM Ub-Rhod was used. Ub-Rhod was added to USP16 using a multichannel pipette to a final volume of 10 µL, and measurements were taken immediately every 30 s for 1 h. For the Ub^R3^ kinetics assay, Ub^R3^ was added during enzyme dilution steps to a final concentration of 20 µM, while an equivalent volume of buffer was used in the control. For Michaelis-Menten determination of K_M_ values for the USP panel, each tested USP was ran at the concentration indicated in **Supplementary Table 1**, while a top concentration of 5 µM of Ub-Rhod was used together with an equimolar concentration and titration of sequester domain (SD) for USP15, USP16 and USP21. Initial data preparation was performed in MARS data analysis software prior to subsequent kinetics analysis. To convert fluorescence intensity for kinetics, raw slopes (FI/min) were divided by the fluorescence intensity range (Max FI – Min FI) obtained from the standard. A linear regression from T = 0 to T = 3 min was then used to determine initial rates, which were subsequently adjusted for enzyme concentration and converted to s⁻¹. Turnover rates (s⁻¹) were plotted against substrate concentration and fit using a non-linear regression Michaelis– Menten model. Data extraction and unit conversion were performed in Excel, and kinetics calculations and graphs were prepared in PRISM (GraphPad).

### Product inhibition and sequestration assays

USP16 and its variants were run at 4 nM and Ub-Rhod at 1 µM. Wild-type Ub or the SD was added during the enzyme dilution steps to a final concentration of 2 µM, while an equivalent volume of buffer was used in the control. For domain-specific variant assays, Ub^R3^, Ub^G76V^, and Ub^Dud^ were added at 500 µM in the same manner, and measurements were taken immediately every 30 s for 1 h. For the USP panel, concentrations of enzyme and Ub are indicated in **Supplementary Table 1**, with the SD included at 5 µM.

### Software & Data Analysis

Sequence alignments and Phylogenetic tree analysis were performed and exported through Jalview^100^ using the ClustalOmega online-tool. Figures were prepared in Adobe Illustrator.

### Data availability statement

Constructs and reagents are available from the corresponding author and Addgene. Coordinates and crystallographic structure factors for USP16 ZnF-UBP and USP49 ZnF-UBP have been deposited with the protein data bank under accession codes 9YCW (USP16 ZnF-UBP) 9YC2 (USP49 ZnF-UBP). Uncropped versions of all gels and blots are provided in **Supplementary Information**.

---

**Extended Data Fig. 1.**
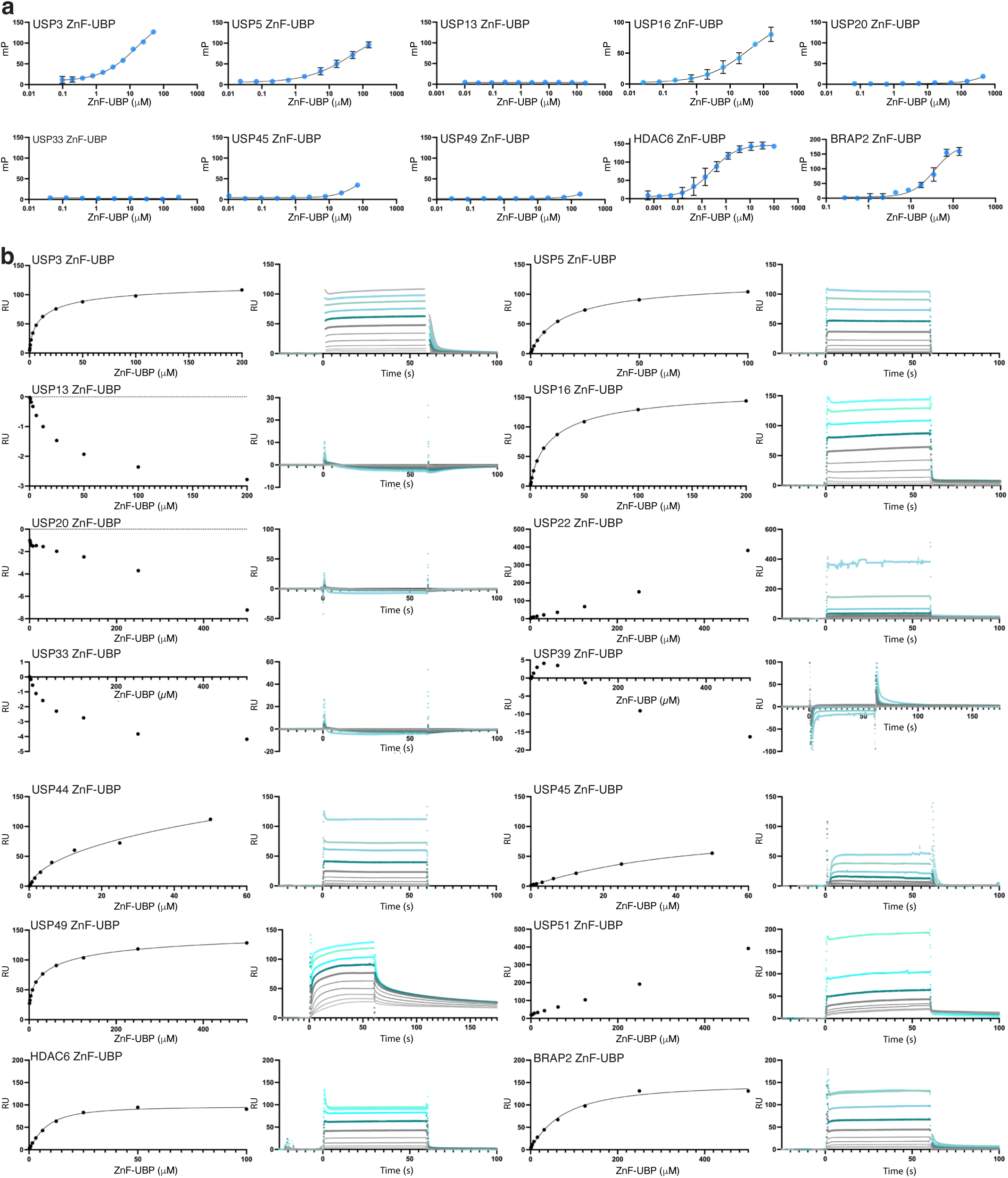
Znf-UBP ubiquitin binding studies. **a,** ZnF-UBP binding against a FITC-labelled Ub C-terminal peptide (FITC-Ahx-RLRGG) measured using Fluorescence Polarization (FP). Indicated ZnF-UBP domains were of sufficiently high yield to be assessed, yet only HDAC6 and BRAP2 yielded full binding curves. Normalised data plotted from *n =* 3 experiments, except for USP45 (*n =* 2). Error bars represent standard error from the mean. **b,** Surface plasmon resonance (SPR) binding curves (*left*) and sensorgrams (*right*) for ZnF-UBP:Ub interactions, with Ub immobilised via amine-coupling. Representative experiments are shown from *n =* 3 independent repeats, except for USP44, USP45 and USP51 (*n =* 2).

**Extended Data Fig. 2.**
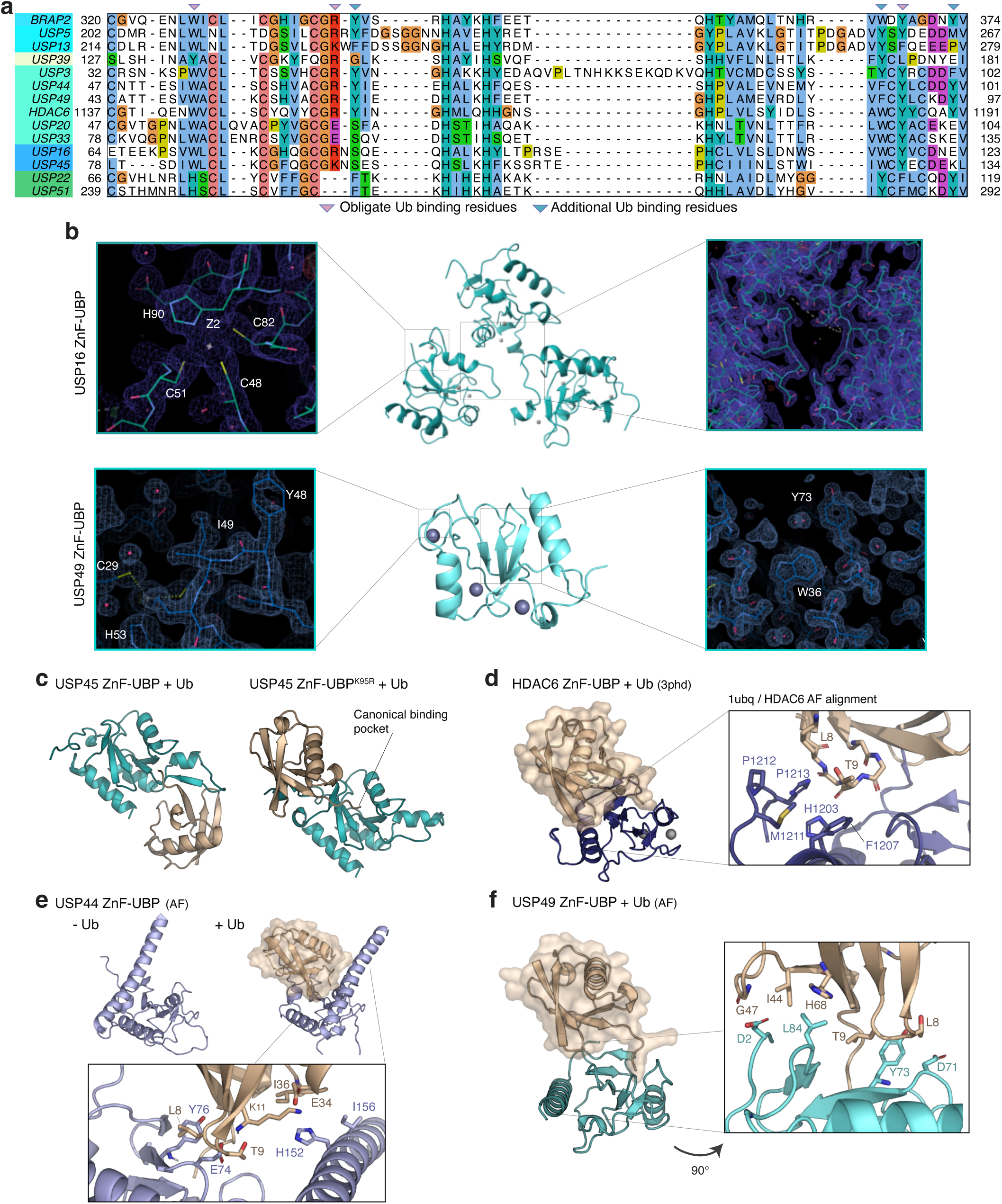
Structural details of the ZnF-UBP domains. **a,** Sequence alignment of ZnF-UBP domains highlighting known and predicted Ub binding residues. **b,** (*Centre*), the asymmetric units of the USP16 (*top*) and USP49 (*bottom*) ZnF-UBP crystal structures. Zn ions are indicated as grey spheres. Close-ups show 2|F_O_|-|F_C_| electron density maps contoured at 1σ, focussing on areas of interest, including density for the Ramachandran outlier Ile49 in USP49 ZnF-UBP. **c,** AlphaFold3 ZnF-UBP:Ub models for wild-type USP45 ZnF-UBP (consistently predicting a non-canonical binding event) vs a K95R mutation (predicting a canonical binding event involving the UbCt pocket). **d,** HDAC6 ZnF-UBP:Ub co-structure (pdb-id 3phd)^16^ aligned with its AlphaFold3 model and Ub (pdb-id 1ubq). HDAC6 binds the Ub TEK box. **e,** AlphaFold3-modelling of apo USP44 ZnF-UBP, in which a preceding α-helix blocks the UbCt pocket. When modelled with Ub, the helix moves to enable UbCt binding and secondary interactions with the Ile36 patch on Ub. **f,** AlphaFold3-predicted model of USP49 bound to Ub showing secondary interactions with the TEK box and Ile44 patch.

**Extended Data Fig. 3.**
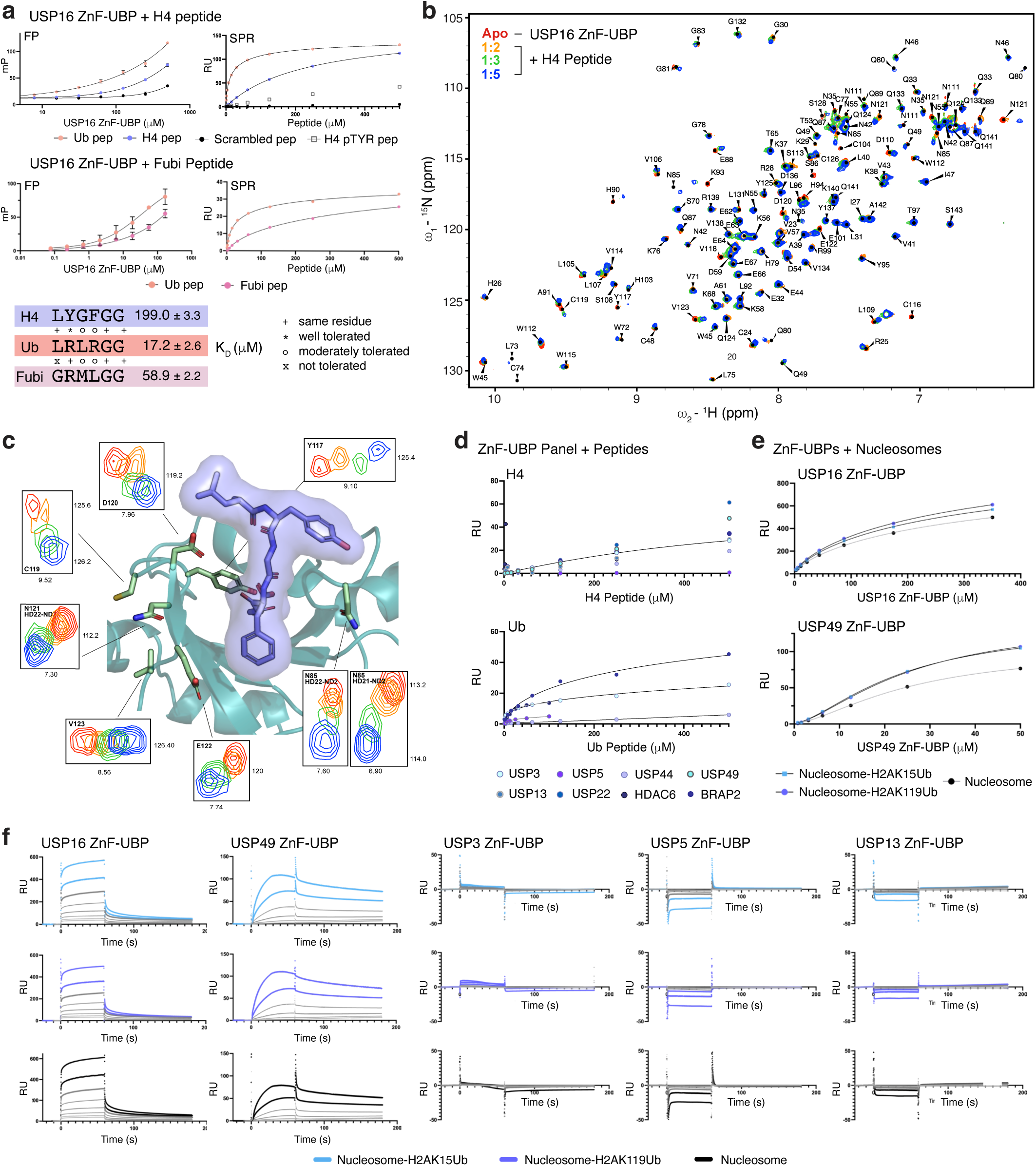
**a,** Binding *(top*) and sequence (*bottom*) comparison of histone H4 and Fubi peptides to UbCt peptide. Binding experiments by FP (left, mean values and s.e.m from *n =* 3 experiments) and SPR (right, *n =* 3 experiments). For SPR, the USP16 ZnF-UBP domain was immobilised by amine coupling. Sequences are compared with residue-tolerance annotations^51^. K_D_ values are derived from SPR experiments, with errors representing s.d. from the mean. H4 pTYR is a phosphorylated (Y99) variant of the H4 peptide testing the effect of prospective histone tail PTMs on ZnF-UBP binding. **b,** Overlay of the 15N-HSQC spectra of free USP16 ZnF-UBP (Apo, in red) with an increasing ratio of histone H4 peptide across three titrations (orange, green and blue), as indicated over the spectra. Peaks were assigned based on the previous assignments available^22^. **c.** Zoom-ins of notable resonance shifts highlighted on AlphaFold3 model of histone H4 peptide-bound USP16 ZnF-UBP. **d,** SPR experiment with immobilised ZnF-UBP domains (amine coupling) to measure binding of histone H4 and UbCt peptides. Representative data from *n =* 2 experiments. Only USP3 (fitted) demonstrated formation of a binding curve against histone H4. **d,** USP16 ZnF-UBP and USP49 ZnF-UBP binding against a panel of immobilised nucleosome variants. Binding curves derived from sensorgrams in **e.** Vastly different binding kinetics differentiate ZnF-UBP interactions, and in case of USP49 lead to a 10-fold higher affinity due to a slow on-and even slower off-rate. **e.** SPR Sensorgrams for ZnF-UBP domains tested against the nucleosome panel (representative from *n =* 2 SPR experiments).

**Extended Data Fig. 4.**
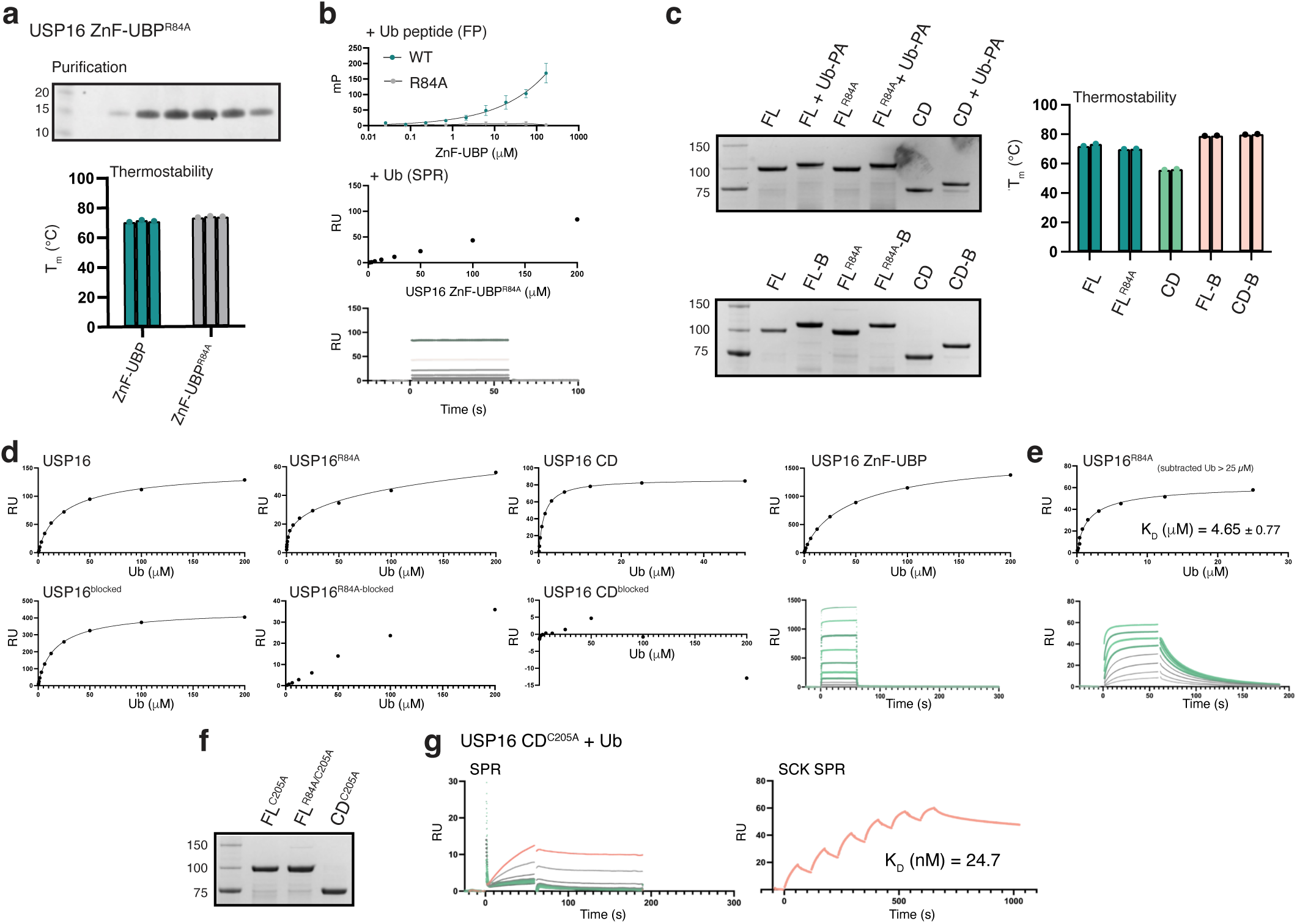
Assessment of USP16’s interaction with Ub. **a,** Coomassie stained SDS-PAGE gel of USP16 ZnF-UBP^R84A^ purification and thermostability profile. **b,** (*top*) UbCt peptide binding to USP16 ZnF-UBP^R84A^ assessed by FP (mean ± s.e.m. from *n =* 3 experiments) and Ub binding by SPR (representative data from *n =* 3 experiments with ZnF-UBP domains amine-coupled to sensor chip). Residual Ub binding is observed. **c,** Coomassie stained SDS-PAGE gel and thermostability profiles of full-length USP16 variants. The top gel shows an activity test, while the gel below shows USP16 variants after purification. Incubation with Ub-PA activity-based probe leads to slower migration due to covalent modification of the catalytic Cys with the Ub probe. FL, full-length constructs; CD, catalytic domain; B, blocked (i.e. modified with Ub-PA). **d,** Ub binding to the USP16 panel including USP16 ZnF-UBP. Unmodified USP16 variants were immobilised via biotinylated AviTag, while Ub-PA, ‘blocked’ variants were immobilised via amine coupling. Representative SPR curves from *n =* 3 experiments. **e,** USP16^R84A^ SPR curves (as in **Fig. 4c**) reanalysed with the highest Ub concentrations (200, 100, 50 µM) removed. The fitted affinity curve shows tighter binding, between wild-type (∼20 µM in ZnF-UBP binding site) and CD (1.2 µM, enzymatic S1 site). Errors represent s.d. from the mean from n=3 experiments. **f,** Coomassie stained SDS-PAGE gel of USP16 variants with a C205A catalytic Cys mutation. **g,** SPR sensorgram and single-cycle kinetics curve for USP16 CD^C205A^. USP16 variants were immobilised via their biotinylated AviTag. Representative SPR data from *n =* 3 experiments. Catalytic Cys to Ala mutation creates catalytic domain with 24 nM Ub affinity.

**Extended Data Fig. 5:**
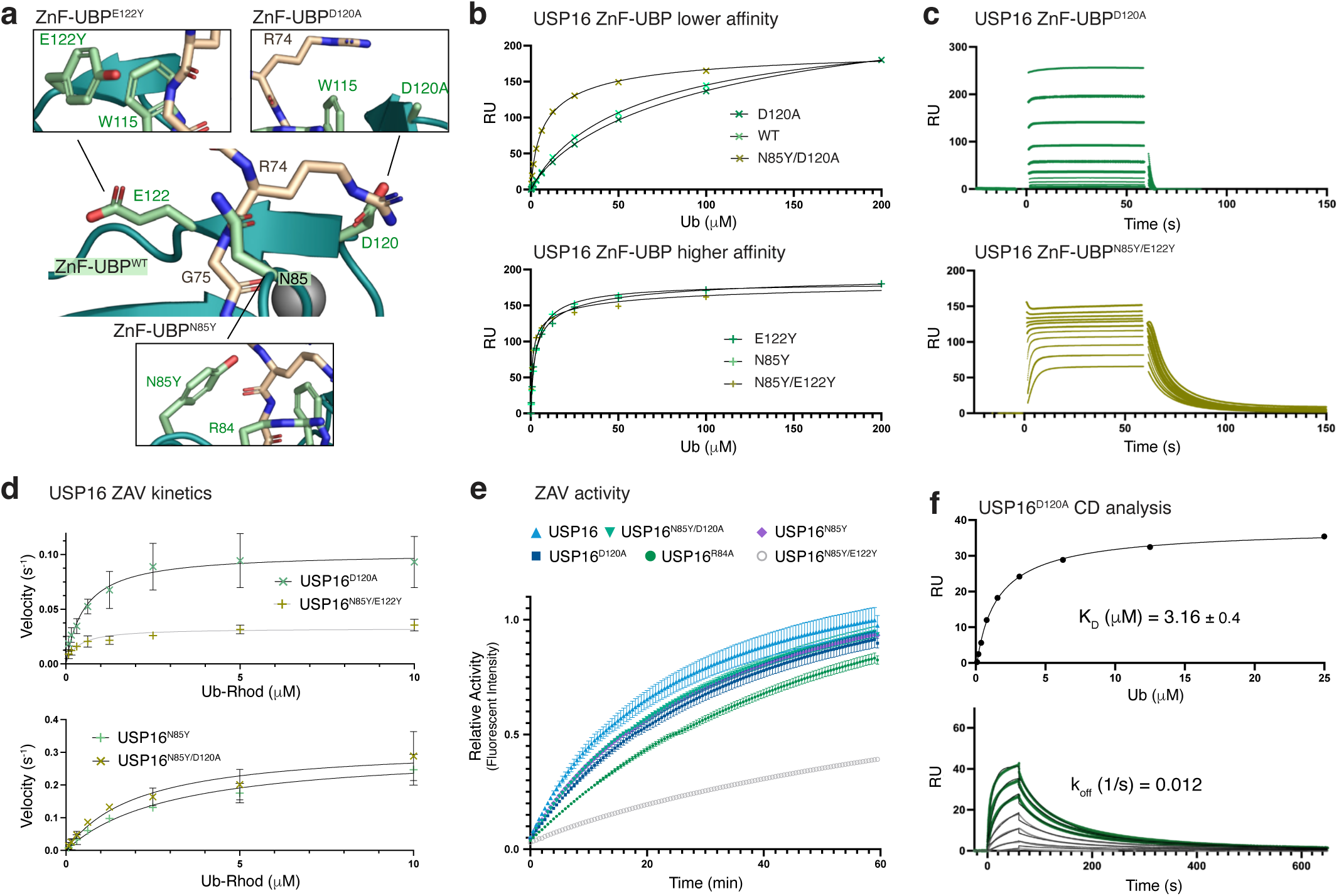
Development of USP16 ZnF-UBP affinity-variants. **a,** UbCt peptide interacting residues mutated in the ZAV panel (**Fig. 5a**) are depicted in the AlphaFold3 model of USP16 ZnF-UBP bound to Ub. **b,** Ub binding affinity variation across the USP16 ZnF-UBP ZAV panel; each ZAV species was amine-coupled to the sensor chip. Representative curves are shown from *n =* 3 SPR experiments. **c,** Exemplary sensorgrams for USP16 ZnF-UBP^D120A^ and USP16 ZnF-UBP^N85Y/E122Y^. **d,** Michaelis–Menten kinetics of full-length USP16 ZAV Ub-Rhod cleavage. Mean ± s.e.m. plotted from *n =* 3 experiments. **e.** Relative activity of all full-length USP16 ZAVs with mean ± s.e.m. taken from *n =* 3 experiments. **f.** Binding kinetics and affinity determination of the catalytic domain of USP16^D120A^. Highest Ub concentrations were excluded to isolate the contribution from the CD and fit using a 1:1 binding model.

**Extended Data Fig. 6.**
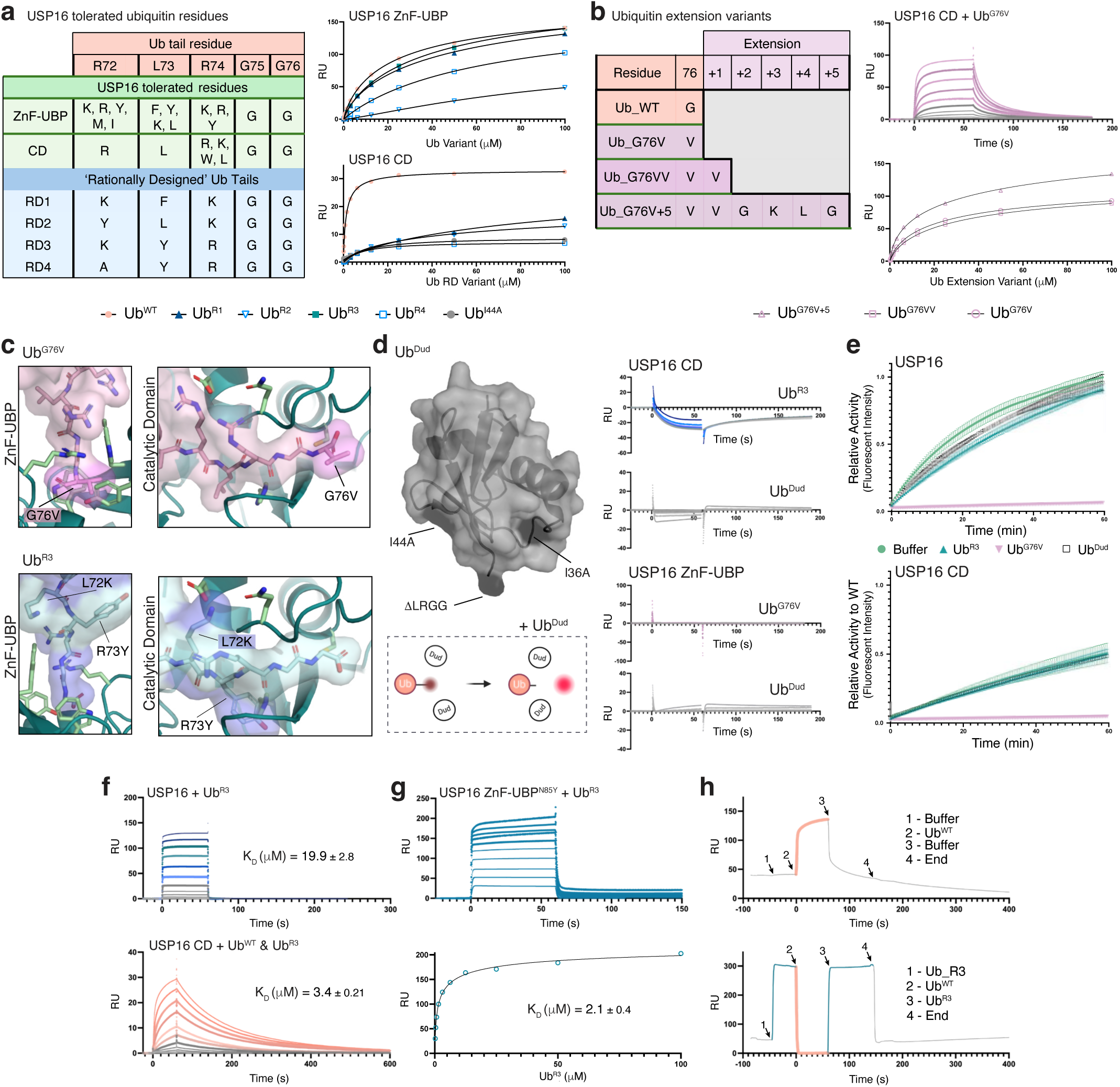
Determining USP16’s domain interdependence by engineering Ub variants. **a,** Residues tolerated by either USP16’s ZnF-UBP domain or USP catalytic domains^51,101^. Four Rational Design (RD) Ub-tail sequences were tested across both domains using SPR (representative curves from *n =* 3 experiments). **b,** Sequences of extended Ub species^102^ binding to USP16 CD (representative sensorgram and curves from *n =* 3 experiments). **c,** Structural overlay of Ub^R3^ and Ub^G76V^ across the ZnF-UBP and CD with mutations labelled. **d,** Structural representation of Ub^Dud^ and assay schematic (left). Comparison of Ub^Dud^ binding to Ub^R3^ and Ub^G76V^ (*right*, representative SPR data from *n =* 3 experiments). **e,** Relative activity of USP16 and USP16 CD Ub-Rhod cleavage supplemented with Ub variants (mean plotted from *n =* 3 ± s.e.m). **f,** Sensorgrams of Ub^R3^ binding full-length USP16 (*top*) and standard USP16 CD Ub binding SPR experiment supplemented with 200 µM Ub^R3^ (*bottom*) (representative data from *n =* 2 experiments). **g,** Binding affinity determination of Ub^R3^ against USP16 ZnF-UBP^N85Y^. Representative data from *n =* 2 SPR experiments. **h,** SPR ABA control experiments for Ub only/unmixed ABA injections (representative experiment from *n =* 2 experiments).

**Extended Data Fig. 7.**
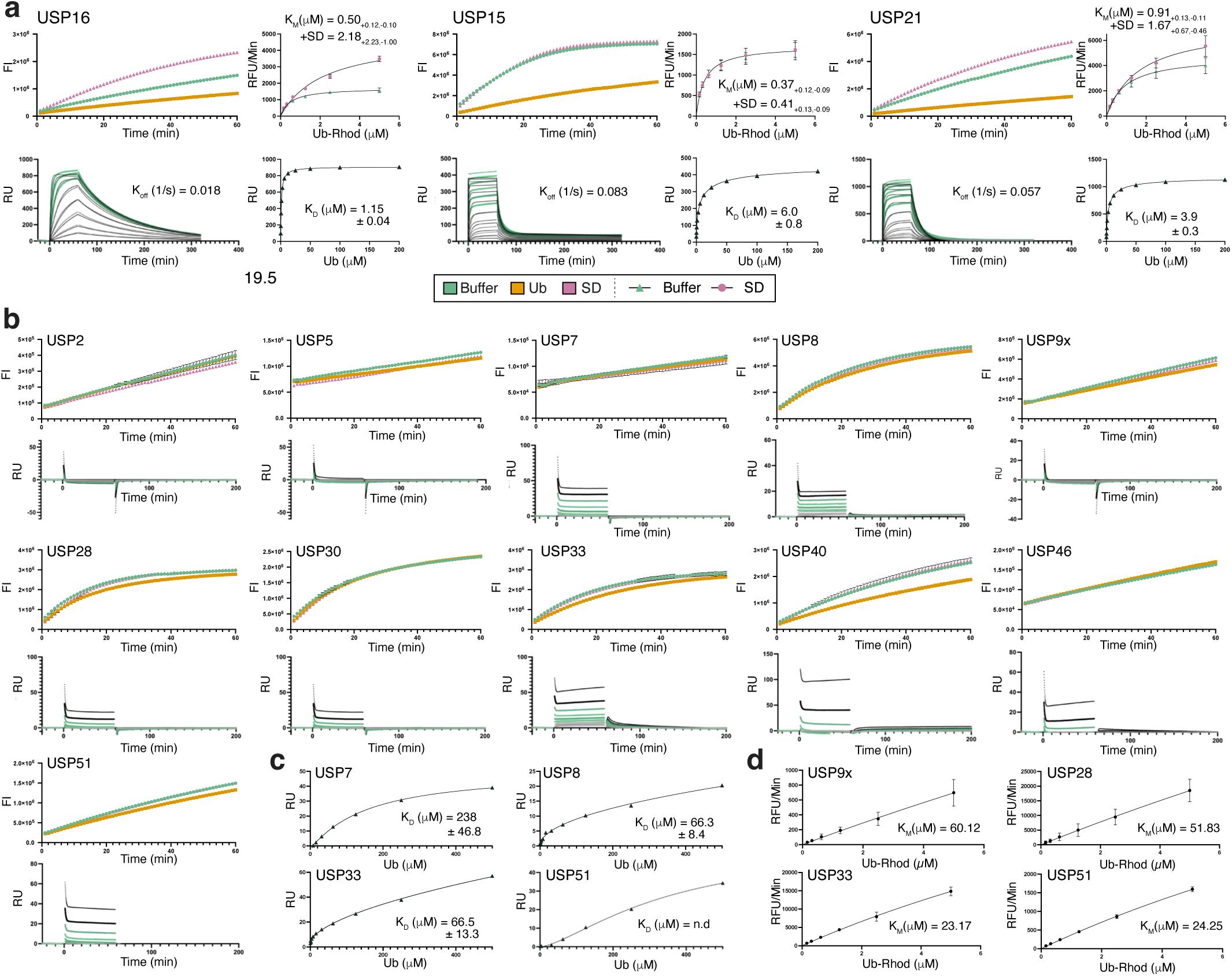
Determination of Ub kinetic and binding properties of the USP panel. **a,** Ub binding and kinetic profiles for indicated SD-affected species. *top left*: representative raw relative activity data (n = 3 experiments); *top right*: Michaelis–Menten kinetics of Ub-Rhod cleavage ± SD domain (mean ± s.e.m., n = 2); *bottom*: representative SPR sensorgram with 1:1 binding model fit and corresponding affinity curve (representative from n=2). **b,** Raw relative activity data (*top*) and SPR binding sensorgrams (*bottom*) for each USP CD tested that were ‘unaffected’ by the SD. Representative data taken from n = 3 (relative activity) and n = 2 (SPR) experiments. **c,** SPR affinity curves for species that were amenable to affinity analysis. Representative curve taken from n = 2 experiments. **d.** Michaelis– Menten kinetics of Ub-Rhod cleavage for selected species (mean ± s.e.m. from n *=* 2 experiments).

**Extended Data Fig. 8.**
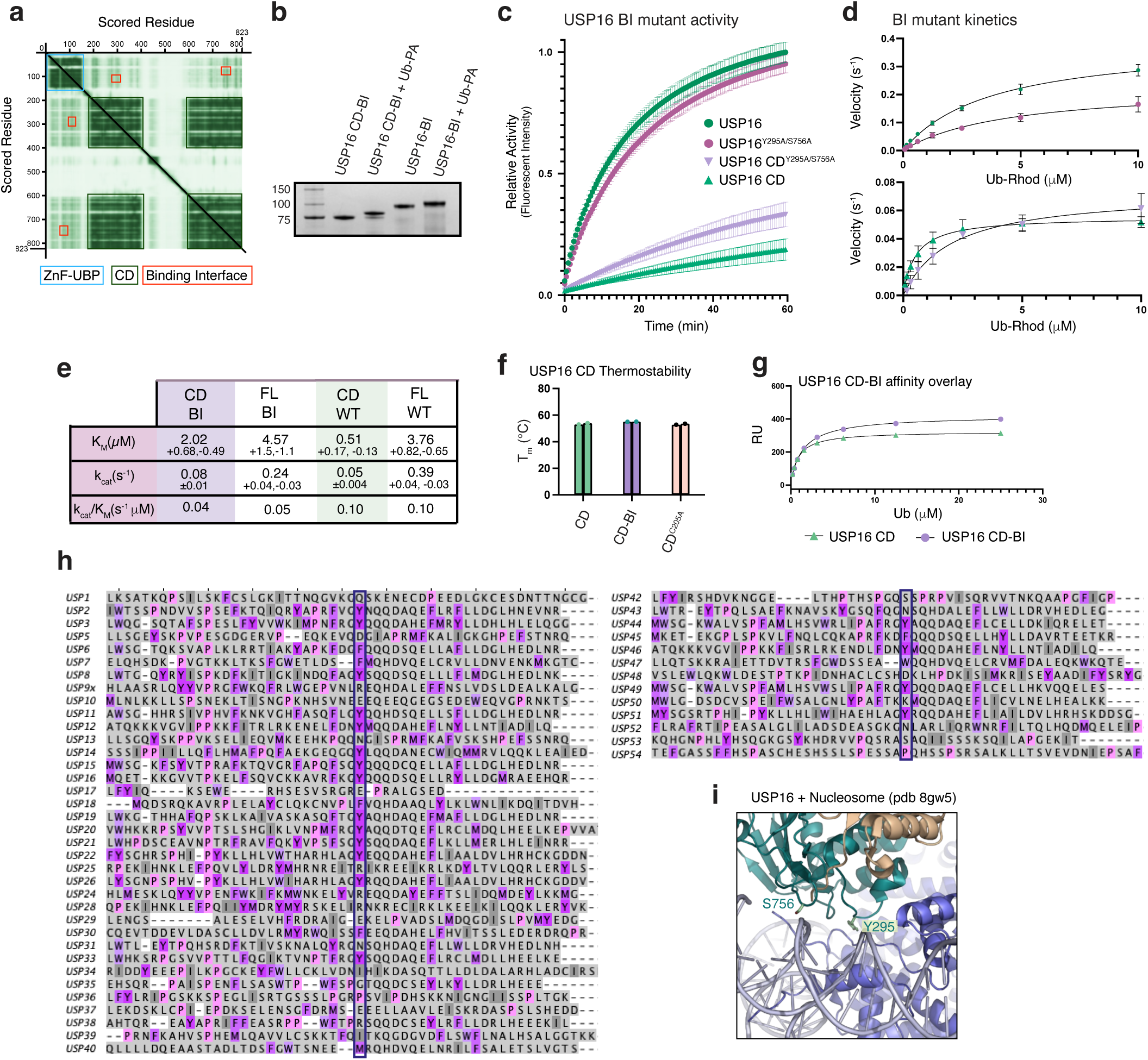
Determination of USP16 Binding Interface validity. **a,** PAE for USP16 model with domains and binding interfaces labelled. Expected position error decreases with intensity of green shade. **b,** Coomassie-stain and Ub-PA activity assay for USP16 BI variants. **c,** Relative activity of Ub-Rhod cleavage for the USP16 BI panel. Plotted Mean ± s.e.m. from *n =* 3 experiments. **d,** Michaelis–Menten kinetics of USP16 BI mutants, determined from Ub-Rhod cleavage assays. Mean ± s.e.m. plotted from *n =* 3 experiments. **e,** Summary of kinetic parameters determined from the Ub-Rhod assays. **f,** Thermostability profiles of USP16 CD variants. **g,** SPR affinity curves for USP16 CD WT and BI variants. Variants were amine-coupled to sensor chips with representative data shown from *n =* 3 experiments. **h,** Sequence-alignment of USP catalytic domains highlighting well-conserved Tyr residue. **i,** USP16:nucleosome binding interface from the USP16:nucleosomeH2AK119Ub co-structure (pdb-id 8wg5)^35^.

**Extended Data Table 1.** Data collection and refinement statistics (molecular replacement).

|  | USP16 ZnF-UBP | USP49 ZnF-UBP |
| --- | --- | --- |
| **Data collection** |  |  |
| Space group | $P3_1$ | $P2_12_12_1$ |
| Cell dimensions |  |  |
| $a, b, c$ (Å) | 84.39, 84.39, 50.43 | 35.74, 47.22, 55.25 |
| $\alpha, \beta, \gamma$ (°) | 90, 90, 120 | 90, 90, 90 |
| Resolution (Å) | 41.51-1.80 (1.84-1.80) * | 47.22-1.40 (1.42-1.40) |
| $R_{\text{merge}}$ | 0.055 (0.661) | 0.065 (0.923) |
| $I / \sigma I$ | 11.5 (1.7) | 11.2 (1.7) |
| Completeness (%) | 99.5 (99.9) | 100 (99.8) |
| Redundancy | 3.3 (3.4) | 5.0 (5.1) |
| **Refinement** |  |  |
| Resolution (Å) | 41.51 – 1.80 | 35.9 – 1.4 |
| No. reflections | 120904 (37022) | 96213 (18997) |
| $R_{\text{work}} / R_{\text{free}}$ | 0.1298/0.1633 | 0.1534/0.1965 |
| No. atoms |  |  |
| Protein | 2615 | 887 |
| Ligand/ion | 27 | 19 |
| Water | 284 | 153 |
| $B$-factors | | |
| Protein | 31.6 | 21.7 |
| Ligand/ion | 34.1 | 24.4 |
| Water | 35.5 | 37.7 |
| R.m.s. deviations |  |  |
| Bond lengths (Å) | 0.010 | 0.005 |
| Bond angles (°) | 0.774 | 0.833 |

A single crystal was used for each data collection. \*Values in parentheses are for highest-resolution shell.

