## Supplementary Information for "ZnF-UBP domains regulate deubiquitinase activity by relieving ubiquitin product inhibition"

Jack A. Alexandrovics[1], Rashmi Agrata[1], Philipp Schenk[1], Anthony Cerra[1], Ueli Nachbur[1], Jeffrey J. Babon[1] and David Komander[1,*]

[1] Walter and Eliza Hall Institute of Medical Research, 1G Royal Parade, Parkville, VIC 3052, Australia; and Department of Medical Biology, University of Melbourne, Melbourne, VIC 3000, Australia.

### Table of Contents

Supplementary Text 3

Supplementary Table 1: USP panel details 5

Supplementary Table 2: ZnF-UBP domain constructs and purification details 6

Supplementary Table 3: USP16 variant constructs and purification details 7

Supplementary Information: Uncropped gels 8

**Supplementary Text**

**Further insights from ZnF-UBP structural modelling**

The USP5 ZnF-UBP bound Ub in a specific orientation influenced by secondary interactions between the Ile36 patch on Ub and a loop on the ZnF-UBP domain (**Fig. 1b**). In contrast to USP5, the different binding orientation of Ub with HDAC6 (**Fig. 1b**) was suggested to be exclusively due to flexibility in Ub's C-terminus[16]. Density was missing in key areas of the reported structure (pdb-id 3phd), and both Alphafold3 modelling and structural alignment of Ub (pdb-id 1ubq) revealed a putative binding interface between Ub's TEK box and a hydrophobic patch on HDAC6[71] (**Extended Data Fig. 2d**). We observed similar orientations and surfaces interactions predicted in related ZnF-UBPs USP44 and USP49. Notably, in AlphaFold3-models of USP44, a prominent $\alpha$-helical extension is predicted to span across the UbCt binding pocket. In modelling USP44 with Ub, the helix shifts to free up Ub access to the UbCt pocket, and shows auxiliary interactions with the Ub TEK box and Ile36 hydrophobic patches (**Extended Data Fig. 2e**). Notably in USP49 ZnF-UBP, the discrepancy of strong Ub interaction ($K_D$ = 22.3 µM) versus lack of detectable UbCt peptide binding might also be explained by similar extensive secondary interactions between the ZnF-UBP with the Leu8 loop and the Ile44 patch of Ub (**Extended Data Fig. 2f**). These predictions require experimental validation.

**Generating specific Ub variants**

Three variations of Ub were engineered to specifically probe the function of each domain in USP16. For a ZnF-UBP specific Ub variants, we generated four rational designs (RDs) informed by previous residue-tolerance tests of ZnF-UBP and USP domains[51,76] (**Extended Data Fig. 6a**). A ZnF-UBP-specific variant, R3, incorporated L72K and R73Y mutations in the C-terminal tail. I36A and I44A mutations were introduced to further ablate interactions with the CD. This variant only bound the ZnF-UBP domain, and not the CD (**Fig. 6b**). To generate a CD specific Ub variant, we exploited the necessity of a C-terminal glycine on Ub for ZnF-UBP binding[51] and explored Ub extension-mutants that retain binding to USP domains[77] (**Extended Data Fig. 6b**). The simplest variant, $Ub^{G76V}$, retained CD binding with a similar profile to wild-type Ub, while ablating interaction with the ZnF-UBP (**Fig. 6b, Extended Data Fig. 6b**). Structural modelling indicated that the substitutions in $Ub^{R3}$ and $Ub^{G76V}$ form steric/electrostatic blocks to their non-permitted domains yet preserve accessibility to their intended domains (**Extended Data Fig. 6c**). A non-binding variant was generated, $Ub^{Dud}$, which removed all known interaction surfaces of Ub, and no longer bound either domain (**Fig. 6b, Extended Data Fig. 6d**).

**Supplementary Table 1: USP panel details**

| Species | Boundaries | Assay [c] | Ub [c] (µM) | SPR [c] (µM) | Literature $K_M$ (µM) |
|---|---|---|---|---|---|
| **USP2** | 258-605 | 5 nM | 10 | 5 | 2.4[78] |
| **USP5** | 322-706^795-858 | 10 µM | 10 | 17 | n.a |
| **USP7** | 208-650 | 20 nM | 10 | 5† | 18.9[80] |
| **USP8** | 734-1119 | 2.5 nM | 10 | 5 | 17.3[38] |
| **USP9x** | 1150-1968 | 7.8 nM | 10 | 26 | n.a |
| **USP15** | 248-477^777-942 | 3 nM | 2 | 5† | 0.72[79] |
| **USP21** | 200-565 | 1 nM | 2 | 21 | 0.26[81] |
| **USP28** | 149-703 | 4 nM | 10 | 21† | n.a |
| **USP30** | 54-517 | 10 nM | 10 | 5 | 39.16[38] |
| **USP33** | 179-288^472-515^570-716 | 10 nM | 10 | 21 | n.a |
| **USP40** | 38-320^444-526 | 10 µM | 2 | 22† | n.a |
| **USP46** | FL | 5 µM | 10 | 17 | 30.07[38] |
| **USP51** | SUMO-356-711 | 0.25 nM | 10 | 17 | n.a |

USP2, USP7, USP8, USP15, USP28, USP30, USP33, USP46 were kindly supplied by Prof David Komander origibally from the LMB. USP5, USP21 USP40, USP51 were previously generated by Anthony Cerra and USP9x by Philipp Schenck.

USPs were typically diluted 1:10 Sodium Acetate pH 5.5 for amine coupling. † Indicates pH 4.5 was used.

n.a = not available

**<u>Supplementary Table 2: ZnF-UBP domain constructs and purification details</u>**

| Species | Boundaries | Storage buffer* |
|---|---|---|
| **USP5** | 171 – 289 | 150 mM NaCl, pH 7.5 |
| **USP13** | 188 – 301 | |
| **USP16** | 22 – 144 | 150 mM NaCl, pH 8.0 |
| **USP20** | 4 – 93 | 150 mM NaCl, pH 8.0 |
| **USP22** | 18 – 139 | 10% (v/v) glycerol (Gly), 500 mM NaCl, pH 7.0 |
| **USP33** | 38 – 124 | 150 mM NaCl, pH 8.0 |
| **USP39** | 96 – 199 | 10% (v/v) Gly, 500 mM NaCl, pH 8.2 |
| **USP44** | 1 – 94 | 10% (v/v) Gly, 500 mM NaCl, pH 7.5 |
| **USP45** | 38 – 173 | 10% (v/v) Gly, 500 mM NaCl, pH 8.2 |
| **USP49** | 1 – 103 | 150 mM NaCl, pH 8.0 |
| **USP51** | 190 – 320 | 10% (v/v) Gly, 500 mM NaCl, pH 7.0 |
| **HDAC6** | 1109 – 1209 | 150 mM NaCl, pH 8.0 |
| **BRAP2** | 247 – 383 | 150 mM NaCl, pH 7.5 |

All constructs were prepared in pOPINK vectors with an N-terminal GST-tag and kanamycin selection marker.
*All storage buffers contained 20 mM HEPES & 2 mM TCEP.

#### Supplementary Table 3: USP16 variant constructs and purification details

| Species | Boundaries | Mutation(s) | Purification | Storage buffer* |
|---|---|---|---|---|
| **Full-length** | 1-823 | n.a, † | GST-Affinity, SEC (HiLoad Superdex pg200), IEC (MonoQ 5/50 GL) | 165 mM NaCl, pH 8.0 |
| | | R84A, † | | |
| | | D120A, † | | |
| | | N85Y, †, ‡ | | |
| | | C205A† | | |
| | | R84A/C205A | | |
| | | N85Y/D120A, ‡ | | |
| | | N85Y/E122Y, ‡ | | |
| | | Y295A/S756A‡ | | |
| **Catalytic Domain** | 192-823 | n.a† | GST-Affinity, SEC (HiLoad Superdex pg200) | |
| | | C205A† | | |
| | | Y295A/S756A‡ | | |
| **ZnF-UBP** | 22 – 144 | n.a† | GST-Affinity, SEC (HiLoad Superdex pg75) | 150 mM NaCl, pH 8.0 |
| | | R84A | | |
| | | D120A | | |
| | | N85Y/D120A | | |
| | | E101Y | | |
| | | N85Y | | |
| | | N85Y/E101Y | | |

Most constructs were prepared in pOPINK vectors an N-terminal GST-tag and kanamycin selection marker except where noted. *All storage buffer contained 20 mM HEPES & 1 mM TCEP. † Indicates an N-terminal Avi-tag variant was produced. ‡ Indicates a Vectorbuilder prepared construct in custom pET17b vector.

Fig 1c

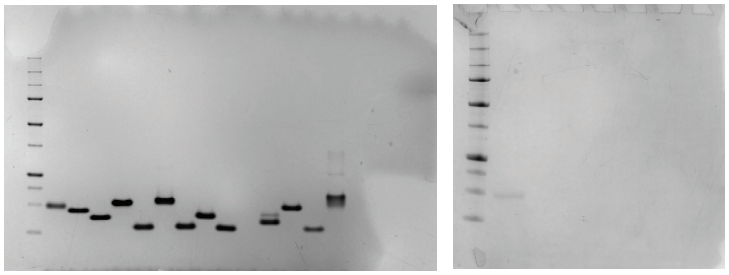

Ext Data Fig 4a

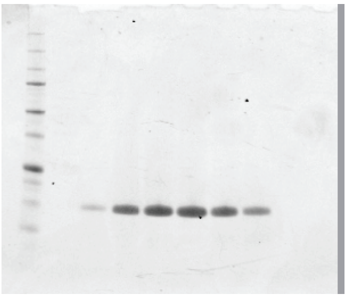

Ext Data Fig 4c (1)

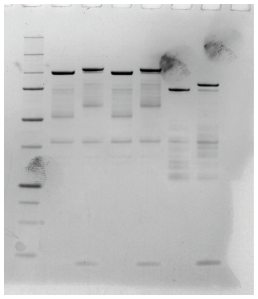

Ext Data Fig 4c (2)

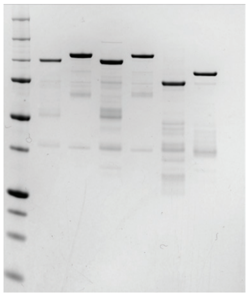

Ext Data Fig 4f

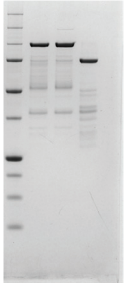

Fig 5a

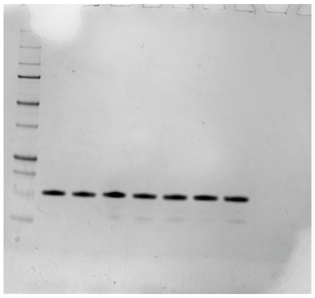

Fig 5b

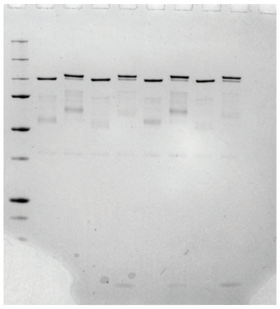

Fig 6b

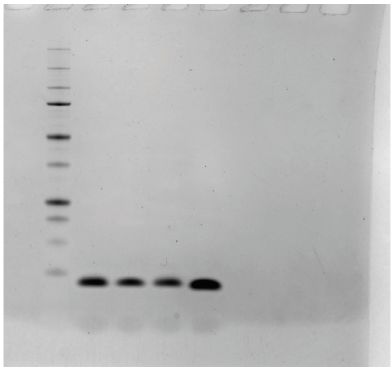

Ext Data Fig 8b

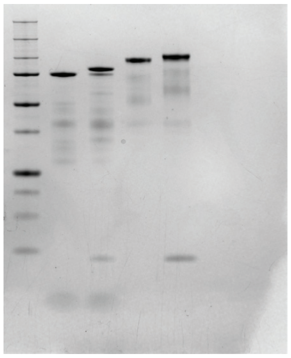

Supplementary Information: Uncropped gels used in this study
